# Development of Degraders and 2-pyridinecarboxyaldehyde (2-PCA) as a recruitment Ligand for FBXO22

**DOI:** 10.1101/2025.08.19.671158

**Authors:** Tian Qiu, Zhe Zhuang, Woong Sub Byun, Zuzanna Kozicka, Kheewoong Baek, Jianing Zhong, Abby M. Thornhill, Julia K. Ryan, Katherine A. Donovan, Eric S. Fischer, Benjamin L. Ebert, Nathanael S. Gray

## Abstract

Targeted protein degradation (TPD) is a promising therapeutic strategy that requires the discovery of small molecules that induce proximity between E3 ubiquitin ligases and proteins of interest. FBXO22 is an E3 ligase that is overexpressed in many cancers and implicated in tumorigenesis. While FBXO22 was previously identified as capable of recognizing ligands bearing a primary amine degron, further investigation and development of recruitment ligands is required to enable its broader utility for TPD. Here, we describe the discovery of chemical probes that can either selectively degrade FBXO22 or recruit this ligase for TPD applications. First, we describe AHPC(Me)-C6-NH_2_ as a potent and selective FBXO22 degrader (DC_50_ = 77 nM, D_max_ = 99%) that is suitable for interrogating the effects of FBXO22 loss of function. Further, we discovered that the simple hexane-1,6-diamine acts as a minimal FBXO22 self-degrader, whereas shorter C4 (putrescine) to C5 (cadaverine) analogs, found in mammalian cells, do not induce degradation. Finally, we found that 2-pyridinecarboxaldehyde (2-PCA) functions as a novel electrophilic degron capable of forming a reversible thioketal with cysteine 326 for recruiting FBXO22. Conjugating 2-PCA to various ligands successfully induced FBXO22-dependent degradation of BRD4 and CDK12. Collectively, these chemical probes will facilitate the study of FBXO22 biology and broaden its applicability in TPD.

## Main

Targeted protein degradation (TPD) has emerged as a compelling alternative to conventional small-molecule inhibition by harnessing the cell’s natural ubiquitin–proteasome system (UPS) to eliminate proteins of interest (POIs)^1-3^. Unlike traditional inhibitors that merely block protein activity, TPD removes the entire protein, thereby abolishing its functions and interactions. TPD primarily employs two types of small molecules: (1) heterobifunctional proteolysis-targeting chimeras (PROTACs), which connect a POI-binding ligand to an E3 ligase recruiter via a suitable linker^4, 5^; and (2) molecular glue degraders (MGDs), which use a monovalent ligand to stabilize interactions between a POI and an E3 ligase^6, 7^. Both strategies promote ternary complex formation, polyubiquitination, and subsequent proteasome-mediated degradation. Despite the growing repertoire of degradable proteins, most TPD approaches still rely on recruiting either cereblon (CRBN)^8^ or von Hippel–Lindau (VHL)^9^ due to the availability of well-described ligands for these ligases. This overreliance presents several challenges, including suboptimal degradation of certain proteins due to incompatible surface topologies, limited expression of CRBN or VHL in some cell types, and the resistance induced by reduced expression of the E3^10-12^. These limitations underscore the need to identify and validate additional ligandable E3 ligases to expand the therapeutic potential of TPD.

The UPS is a central regulator of protein homeostasis and a validated therapeutic target, as demonstrated by the clinical success of proteasome inhibitors in oncology. However, these agents often exert broad effects leading to proteotoxic stress, highlighting the need for more selective approaches^13^. Targeting E3 ubiquitin ligases—the components that confer substrate specificity within the UPS—offers a more precise strategy. E3 ligases play critical roles in diverse cellular processes and have been implicated in cancer, neurodegeneration, and inflammatory diseases^14-16^. Yet, most E3 ligases remain pharmacologically inaccessible due to the lack of well-defined ligandable pockets. To date, the few available E3 ligase inhibitors—including those directed at MDM2^17^, IAPs^18, 19^, VHL^20, 21^, and KEAP1^22^—primarily function by disrupting protein-protein interactions (PPIs) with substrates. Despite progress, there remains a stark gap between the number of known human E3 ligases and the small fraction currently targeted by drug-like molecules. Recently, small-molecule degraders of E3 ligases have emerged as a promising alternative modality^23^. Still, their broader application is limited by the scarcity of validated ligands for most E3 ligases. These challenges underscore the need to develop both ligands and degraders targeting therapeutically relevant E3 ligases.

FBXO22 (F-box only protein 22) exemplifies this opportunity and has recently emerged as a promising E3 ligase for TPD applications^24-28^. As a member of the F-box protein family, FBXO22 functions as the substrate receptor for a SCF (SKP1–CUL1–F-box protein) E3 ubiquitin ligase complex. FBXO22 has been implicated in the development and progression of multiple cancers, including liver^29^, colorectal^30^, breast cancers^31^ and leukemia^32^. Analysis of The Cancer Genome Atlas (TCGA) data revealed that FBXO22 mRNA levels are significantly upregulated in a range of human tumor tissues compared to corresponding normal tissues^33^. Functionally, FBXO22 regulates key processes such as cell cycle progression, DNA damage response, and chromatin remodeling by targeting tumor suppressors, transcriptional regulators including the cyclin-dependent kinase inhibitor p21^29^, and the histone demethylase KDM4A^34, 35^. A particularly well-characterized substrate is the transcription factor BACH1 (BTB and CNC homology 1), which plays a key role in orchestrating oxidative stress adaptation^32, 36^. Recent structural studies have revealed how FBXO22 recognizes a heme-regulated degron within the C-terminal region of BACH1, establishing a direct link between cellular heme levels and BACH1 stability^37, 38^. Chemical modulation of FBXO22 activity could serve as valuable tool to further dissect its biological functions. In addition, given the elevated expression in cancer cells, FBXO22 represents an attractive E3 ligase for expanding the scope of TPD approaches with tumor-specific activities.

Here, we report the discovery of a selective FBXO22 degrader and the subsequent optimization of an electrophilic warhead for FBXO22-mediated TPD. We serendipitously discovered that a primary amine contaminant generated during the synthesis of a PROTAC induced degradation of FBXO22. Structure-activity relationship studies of this compound identified the primary amine as being the essential degron. We constructed a small library of primary amine-derivatized CRBN- or VHL-based PROTACs, which was screened to identify the VHL-recruiting compound, AHPC(Me)-C6-NH_2_ as the most potent FBXO22 degrader. Mechanistic studies demonstrated that metabolism of the primary amine to the corresponding aldehyde results in formation of the biologically active AHPC-CHO. AHPC-CHO forms a reversible-covalent thioketal with Cys326 of FBXO22 thereby stabilizing a ternary complex with VHL, which results in ubiquitination and subsequent proteosome-mediated degradation of FBXO22. We also discovered that simple diamino or dialdehyde alkanes can result in FBXO22 self-degradation. Lastly, during our efforts to improve self-degrader activity, we identified 2-pyridinecarboxaldehyde (2-PCA) as a novel reactive warhead for recruiting FBXO22 and developed degraders for targets inlcuding BRD4 and CDK12. Collectively, our findings provide a valuable chemical probe for investigating the biological function of FBXO22 and pave the way for more broadly harnessing FBXO22 in TPD.

### E3 ligand tethered with alkyl amine degrades FBXO22

As part of a chemical biology campaign to expand the degradable proteome, we used whole-cell proteomics to analyze the degradation profiles of our in-house PROTAC library. We observed that FBXO22 was the most prominently downregulated protein (P. Value < 0.001, |fold change| > 1.5) in both HEK293T and Kelly cells treated with ZXH-3-118 (Figure 1A, S1B, C). This compound is a byproduct generated during the synthesis of a WDR5 PROTAC molecule, which consists of a CRBN binder thalidomide^39^, a spacer, and a primary alkylamine (Figure S1A). We confirmed FBXO22 degradation in HEK293T, Kelly, and MOLT-4 cells (Figure S2A, B). Notably, this degradation was completely abolished upon depletion of fetal bovine serum (FBS) (Figure S2C), consistent with recent reports that amine oxidases present in FBS catalyze the metabolic conversion of primary alkylamine-containing compounds into their corresponding aldehyde analogs, thereby enabling engagement with FBXO22^24, 25^. Based on these findings and the structural features of ZXH-3-118, we reasoned that an E3 ligase ligand combined with a primary amine group might be the minimal requirement for designing a degrader of FBXO22 (Figure 1A). Accordingly, we constructed a library of compounds containing various E3 ligase ligands (including CRBN, VHL, MDM2^40^, and IAP^41-44^ and primary amine moieties with various linker lengths.

**Figure 1.**
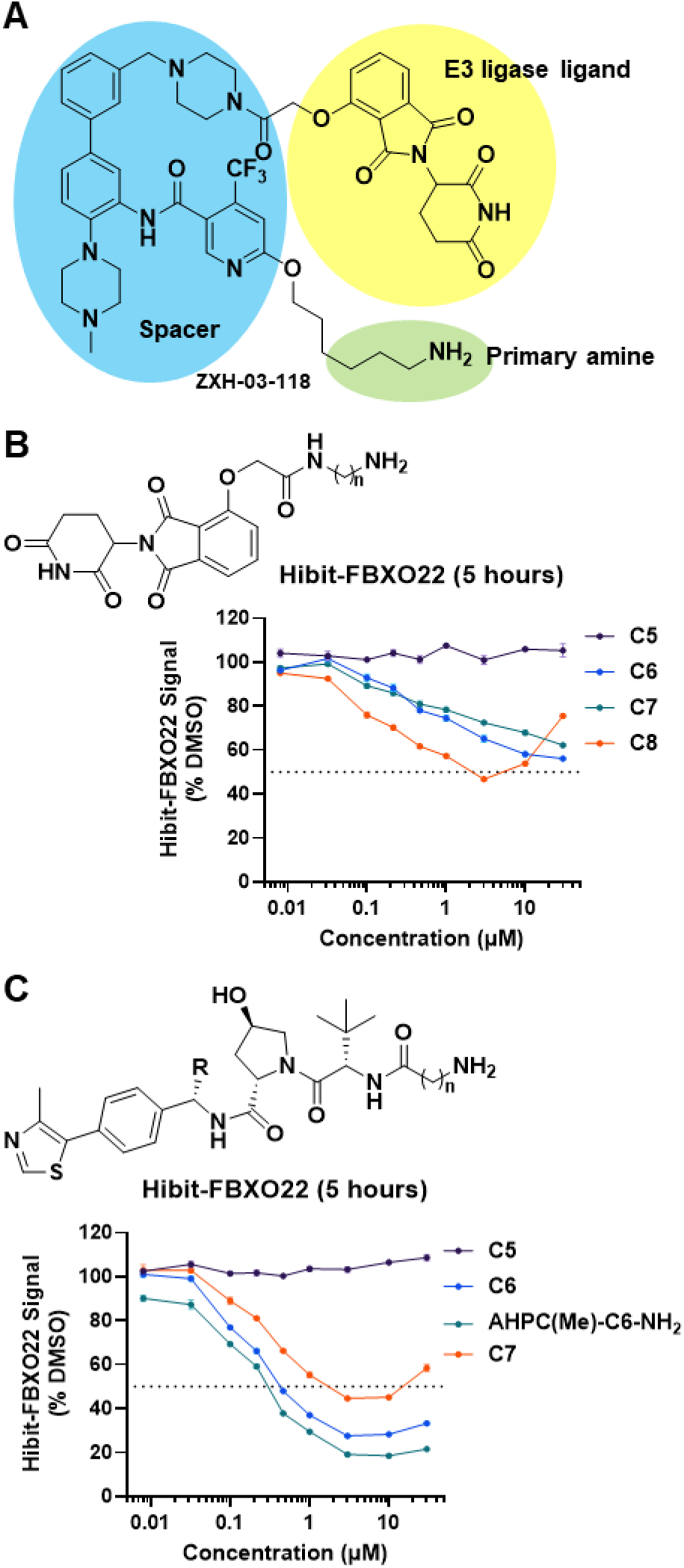
Charting E3 ligand tethered with primary amine for FBXO22 degradation. A) Structure of ZXH-03-118. E3 ligase ligand is highlighted with yellow color, primary amine is highlighted with green color, spacer region is highlighted with blue color. B) General chemical structure of Thalidomide-Amide-Cn-NH_2_ and HiBiT-FBXO22 assay results for Jurkat cells treated with the CRBN ligand-based degraders for 5 hours. C) General chemical structure of AHPC(R)-Cn-NH_2_ and HiBiT-FBXO22 assay results for Jurkat cells treated with the VHL ligand-based degraders for 5 hours.

To facilitate compound screening, we established a HiBiT-FBXO22 assay by stably overexpressing FBXO22 fused to an N-terminal HiBiT-RR tag^45^ in Jurkat cells. The engineered cell lines were treated with compounds across a range of concentrations for either 5 or 18 hours, followed by endpoint measurement of target degradation using a luciferase assay. We initiated the screening with CRBN ligand-based compounds, as the original molecule containing a thalidomide ligand exhibited moderate activity. Simply attaching alkylamine of varying lengths to the 4-position of thalidomide via either O- or NH-linkages did not produce active FBXO22 degraders (Figure S3A, B). Considering FBXO22 is also an E3 ligase, we used Western blot analysis to determine whether compounds could induce CRBN degradation but found no degradation of either protein (Figure S3C). Considering the structural similarity to ZXH-3-118, we hypothesized that incorporating a spacer might enhance molecular flexibility and improve activity. To test this, we prepared a series of thalidomide amido derivatives with different linker lengths. We found that linkers longer than C5 conferred moderate FBXO22 degradation activity, with a C8 linker compound achieving approximately 50% degradation at 3 μM after 5 hours of treatment. In contrast, compounds with PEG linkers remained inactive (Figure 1B, S3D). Extending the treatment duration to 18 hours did not further increase degradation efficacy. These results suggest that thalidomide tethered to primary alkylamine can induce moderate FBXO22 degradation, consistent with our observations for ZXH-3-118.

Next, we examined a series of VHL derivatives based on the AHPC ligand. While linkers shorter than C6 were inactive, the C6 derivative exhibited robust FBXO22 degradation, achieving approximately 70% degradation at 3 μM after 5 hours (Figure 1C, S4). Longer treatment durations further increased the D_max_ to around 80%. Introducing a methyl group at the benzylic position, known to enhance VHL binding affinity, yielded an improved FBXO22 degrader that achieved approximately 80% degradation within 5 hours. We also tested MDM2 and IAP ligand tethered to hexylamine; however, neither induced FBXO22 degradation (Figure S5A, B). Based on these findings, we selected AHPC(Me)-C6-NH_2_ (Figure 2A) as the lead FBXO22 degrader for further investigation.

**Figure 2.**
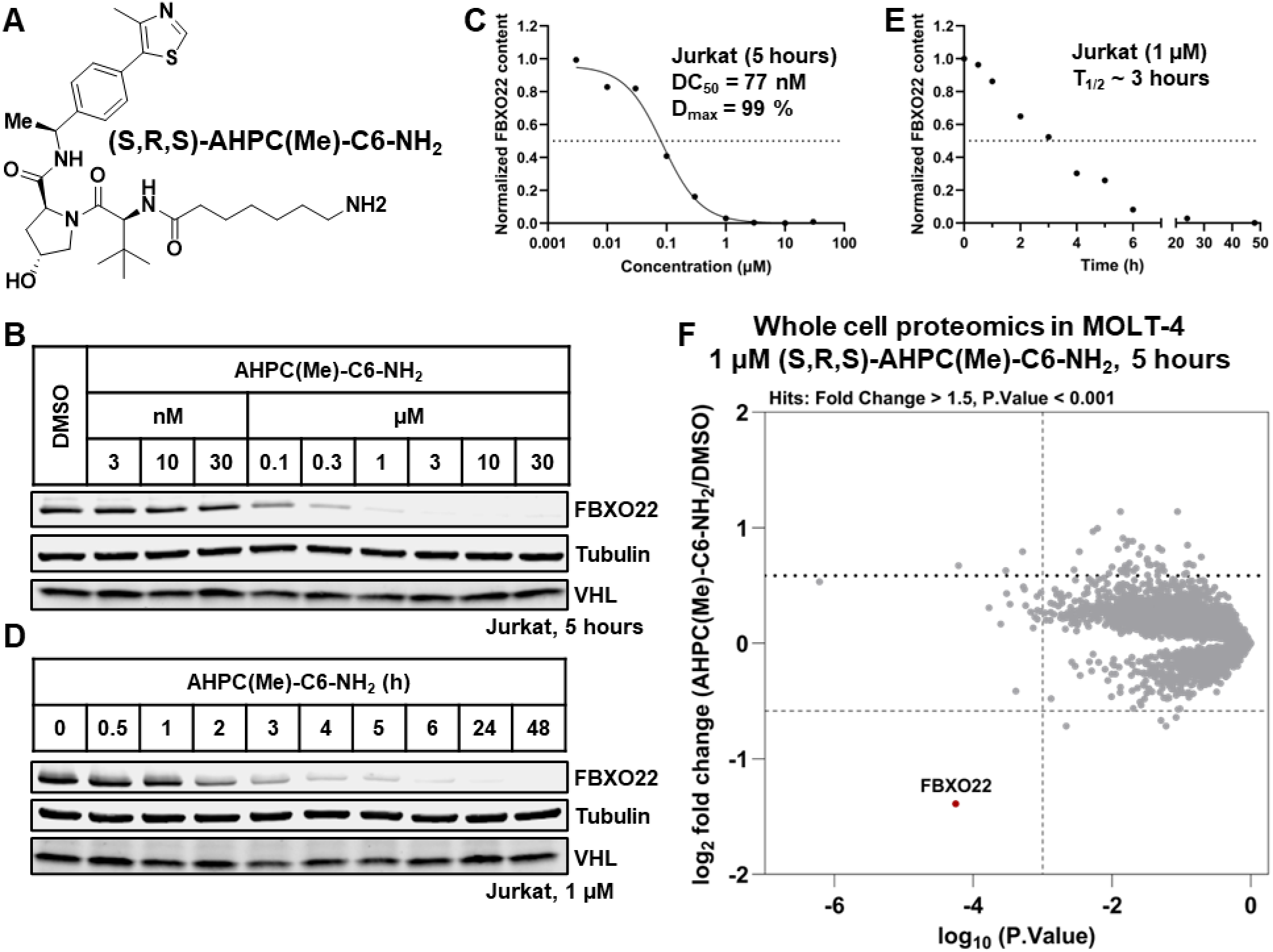
(S,R,S)-AHPC(Me)-C6-NH_2_ is a potent and selective FBXO22 degrader. A) Chemical structure of AHPC(Me)-C6-NH_2_. B) Western blots showing FBXO22 degradation in Jurkat cells treated with the indicated concentration of AHPC(Me)-C6-NH_2_ for 5 hours. C) Quantification of the data shown in B. D) Western blots showing FBXO22 degradation in Jurkat cells treated with 1 μM AHPC(Me)-C6-NH_2_ at indicated time. E) Quantification of the data shown in D. F) Quantitative proteome-wide mass spectrometry in MOLT-4 cells after 5 hours treatment with 1 μM AHPC(Me)-C6-NH_2_.

### AHPC(Me)-C6-NH_2_ is a selective FBXO22 degrader

To evaluate the degradation efficacy of AHPC(Me)-C6-NH_2_ on endogenous FBXO22 protein, we conducted time- and dose-dependent experiments in Jurkat cells. The compound markedly induced the FBXO22 degradation at 1 μM over 5 hours incubation, with a half-maximal degradation concentration (DC_50_) of 77 nM and a D_max_ of 99% (Figure 2B, C). Treatment with 1 μM AHPC(Me)-C6-NH_2_ resulted in FBXO22 degradation beginning as early as after 1 hour, reaching the maximum degradation by 5 hours, and persisted for at least 48 hours (Figure 2D, E). Notably, VHL protein levels remained unaffected, indicating that in the context of FBXO22-VHL PROTACs, FBXO22 is the primary degradable target.

To assess selectivity, we carried out whole-cell proteomics on MOLT-4 cells treated with 1 μM of AHPC(Me)-C6-NH_2_ for 5 hours. Despite its structural simplicity, the degrader exhibited a highly selective degradation profile, with FBXO22 being the only protein efficiently degraded across the entire proteome (Figure 2F). Collectively, these results confirm that AHPC(Me)-C6-NH_2_ is a potent and highly selective chemical degrader of endogenous FBXO22.

### Investigation of the mechanism of AHPC(Me)-C6-NH2 induced FBXO22 degradation

To investigate the mechanism underlying AHPC(Me)-C6-NH_2_ induced FBXO22 degradation, we integrated chemical synthesis and pathway inhibition experiments to assess the roles of the UPS pathway and primary amine metabolic conversion. First, we synthesized two negative control compounds, ZZ7-23-033 and ZZ7-23-034, which feature either a secondary amine (resistant to oxidation to the aldehyde) or an inverted stereocenter (preventing VHL binding), respectively (Figure 3A). Both compounds failed to induce FBXO22 degradation, indicating that both VHL engagement and a primary amine are required for activity (Figure 3B, S6A). Next, pre-treatment with UPS pathway inhibitors—including TAK-243 (E1 inhibitor), MLN4924 (neddylation activating enzyme inhibitor), MG132, and bortezomib (proteasome inhibitors)—blocked FBXO22 degradation. Additionally, pre-incubation with excess VHL ligand (VH032) abolished FBXO22 degradation, confirming the requirement for VHL recruitment. Lastly, we used two conditions to demonstrate the necessity of metabolic activation: (1) pretreatment with aminoguanidine (AG), a pan-amine oxidase inhibitor^46^, blocked FBXO22 degradation in the presence of FBS; and (2) switching from FBS-containing medium to Opti-MEM, which lacks FBS, also prevented degradation (Figure 3C, S6B).

**Figure 3.**
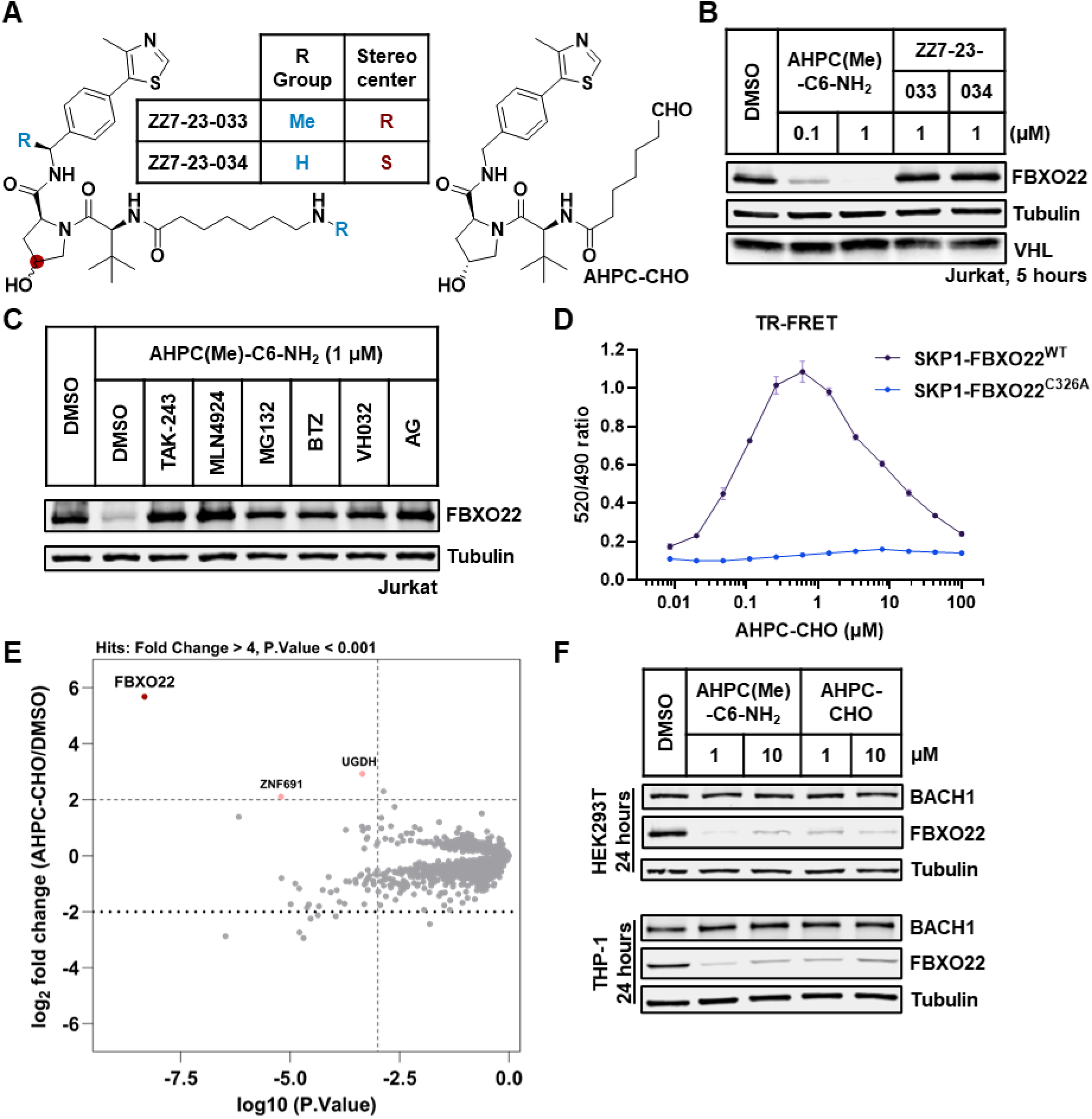
Mechanism of AHPC(Me)-C6-NH_2_ induced FBXO22 degradation. A) Chemical structures of ZZ7-23-033, ZZ7-23-034 and AHPC-CHO. B) Western blots showing FBXO22 degradation in Jurkat cells treated with the indicated compounds for 5 hours. C) Western blots showing FBXO22 degradation in Jurkat cells pre-treated with the indicated inhibitors for 1 hour, followed by treatment with 1 µM AHPC(Me)-C6-NH_2_ for 5 hours. D) TR-FRET assay to measure molecule-dependent ternary complex formation between VCB complex and SKP1/FBXO22 (WT or C326A). Each point represents a duplicate replicates; the mean value line is drawn. E) Scatterplots depicting relative protein abundance following Flag-VHL-EloB/C enrichment from in-lysate treatment with 1 µM AHPC-CHO and recombinant Flag-VHL-EloB/C spike in. Scatterplots display fold change in abundance to DMSO. Significant changes were assessed by moderated t-test as implemented in the limma package with log_2_ FC shown on the y-axis and log_10_ (P.value) on the x-axis. F) Western blots showing protein levels of FBXO22 and its downstream signals in HEK293T and THP-1 cells treated with indicated concentration of AHPC(Me)-C6-NH_2_ and AHPC-CHO for 24 hours.

Based on two previous studies demonstrating that primary amine-containing degraders are metabolized to the corresponding biologically active aldehydes, we synthesized a series of VHL ligand tethered alkyl aldehyde compounds with varying linker lengths (Figure 3A, S7A). In cellular assays, the analog equivalent to AHPC(Me)-C6-NH_2_, namely AHPC-CHO, displayed the highest potency (DC_50_ = 150 nM, D_max_ = 72 %). Analogs lacking a linker (ZZ7-24-008) or with a linker shortened by one carbon (TWQ-07-040) were inactive, indicating a minimal linker length requirement for effective FBXO22 degradation. An analog with a linker extended by one carbon (TWQ-07-041) showed slightly improved efficacy at low concentrations, but the D_max_ only reached 50% (Figure S7B, C). As expected, AHPC-CHO-mediated degradation was blocked by UPS pathway inhibitors, whereas switching to FBS-free Opti-MEM medium had no impact (Figure S7D). In addition, AHPC-CHO outperformed its alkylamine analog at low concentrations but exhibited a more pronounced hook effect—a phenomenon often observed with PROTACs. These findings suggest that conversion of the amine into the aldehyde in FBS-containing medium serves as a prodrug mechanism, maintaining an optimal concentration of the active species to enable sustained FBXO22 degradation.

Building on this validation of AHPC-CHO as an active degrader, we next biochemically investigated its ability to form a ternary complex. To this end, we conducted a TR-FRET assay using four VHL-aldehyde compounds with labeled VHL-EloB/C (VCB complex) and SKP1/FBXO22 (Figure S7E). Most compounds elicited a dose-dependent TR-FRET signal between 10 nM and 1 μM, with a hook effect observed at higher concentrations. In contrast, the linkerless compound ZZ7-24-008 produced negligible TR-FRET signal, only detectable at high concentrations. TWQ-07-040 showed the weakest TR-FRET signal, consistent with its lack of FBXO22 degradation and further supporting that an inadequate linker length impairs effective ternary complex formation. Previous reports suggested that alkyl aldehydes can engage cysteine 326 on FBXO22 to recruit FBXO22 for degradation of FKBP12 and NSD2. We show that introducing a cysteine-to-alanine mutation at this site results in a near-complete loss of ternary complex formation between the VCB complex and SKP1/FBXO22 (Figure 3D). Together, these findings demonstrate that AHPC-CHO facilitates ternary complex formation via cysteine 326 on FBXO22.

Given that whole-cell proteomics indicated high selectivity of AHPC(Me)-C6-NH_2_ for degradation, we next investigated whether this selectivity originated from preferred interaction between VHL and FBXO22 in the presence of degrader. Using immunoprecipitation mass spectrometry (IP-MS)^47^, we incubated MOLT-4 lysates with 1 μM AHPC-CHO and recombinant FLAG-tagged VCB complex. After one hour, the complex was enriched with Anti-FLAG^®^ M2 magnetic beads, followed by label-free quantitative proteomics to identify interactors (Figure 3E). Remarkably, only three proteins were significantly enriched (P. Value < 0.001, log_2_ (fold change) > 2), with FBXO22 being the most enriched protein (P. Value = 4.7 x 10^-9^, log_2_ (fold change) = 5.7) across the whole proteome. These findings support that the observed degradation selectivity arises from the specific recruitment of FBXO22 to the VCB complex.

Having confirmed the potency and selectivity of the AHPC(Me)-C6-NH_2_, we next assessed its activity in additional cell lines. Treatment of THP-1 and HEK293T cells with AHPC(Me)-C6-NH_2_ and AHPC-CHO for 24 hours resulted in robust FBXO22 degradation (Figure 3F). We also probed BACH1, a well-characterized substrate of FBXO22, to gain preliminary insight into downstream effects. In THP-1 cells, we detected a modest increase in BACH1 protein levels, consistent with previous reports using genetic knock-down or knock-out in this cell line^32^. Meanwhile, no clear changes in BACH1 were observed in HEK293T or Jurkat cells. Since there are no direct studies showing that FBXO22 loss affects endogenous BACH1 in these cell lines, we generated FBXO22 knockout Jurkat cell line using CRISPR-Cas9 (Figure S8). Although FBXO22 was efficiently depleted, BACH1 protein levels remained unchanged, suggesting that other E3 ligases may regulate BACH1 stability in this context. Together, these results support the utility of AHPC(Me)-C6-NH_2_ as a chemical biology tool to modulate FBXO22 levels and provide a foundation for future studies on FBXO22 function in diverse cellular environments.

### Discovery of primary diamine as FBXO22 self-degrader

Symmetric homodimerizing PROTACs have been successfully developed for CRBN^48^ and VHL^23^. We investigated whether a primary diamine could induce self-degradation of FBXO22 (Figure 4A). To this end, we screened a series of primary diamines with varying linker lengths, along with their corresponding dialdehyde metabolites, using the HiBiT-FBXO22 assay (Figure 4B). Consistent with prior results for VHL- or CRBN-ligands tethered with alkylamines, primary diamines with C3-C5 linkers showed no activity (Figure 4C, S9A, B). Although glutaraldehyde (1,5-pentanedial) demonstrated strong degradation activity in the Hibit-FBXO22 assay (Figure S9C), it also caused cytotoxicity after 18 hours, likely due to its known protein crosslinking properties (Figure S9D). Among the primary diamines tested, hexane-1,6-diamine exhibited the most potent degradation activity with a D_max_ of approximately 50% at 5 hours, whereas its dialdehyde analog showed only marginal activity. Notably, compounds with linker lengths exceeding C7 lost activity dramatically, indicating that optimal linker length is critical for effective FBXO22 self-degradation.

**Figure 4.**
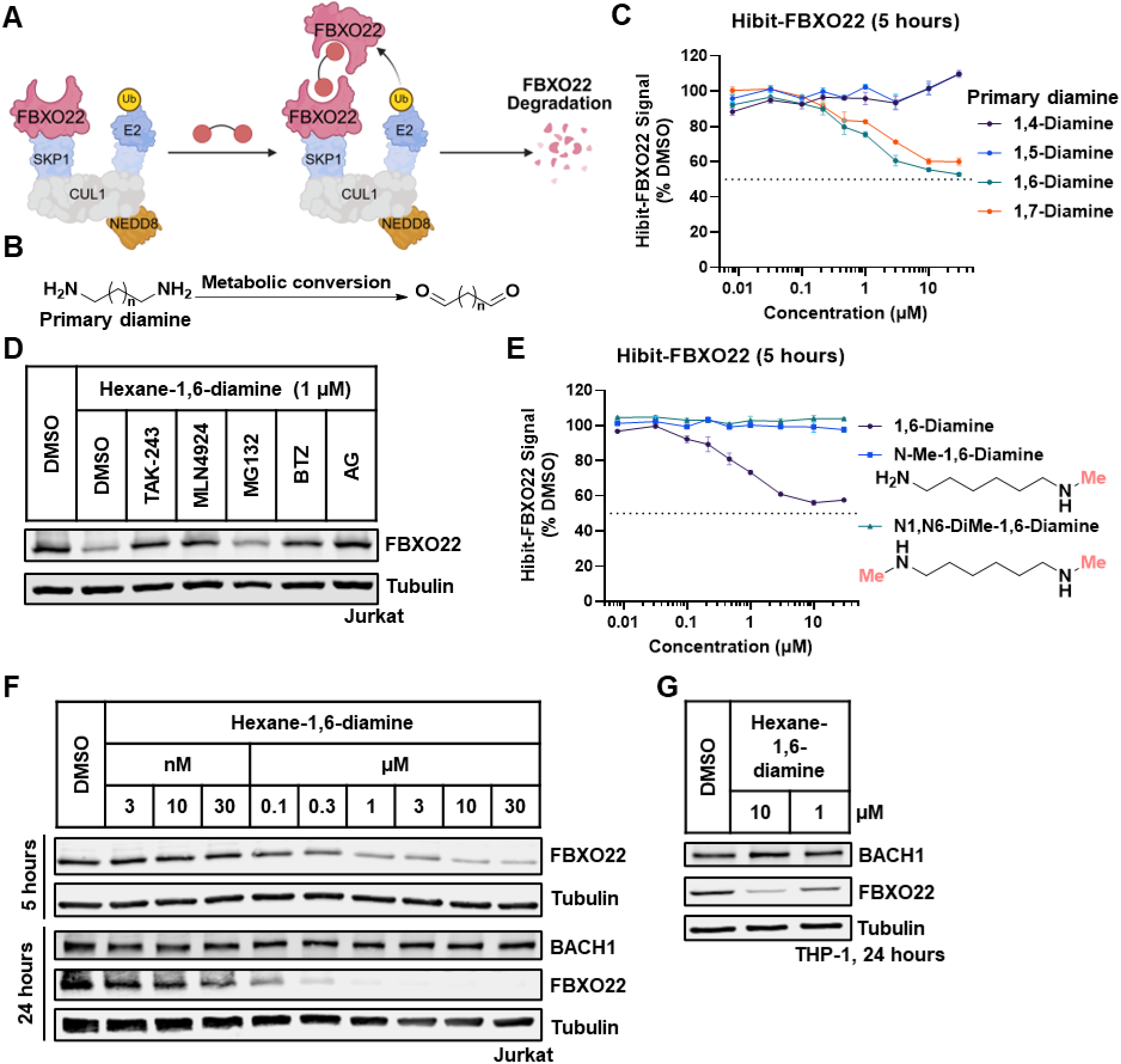
Primary diamine is FBXO22 self-degrader. A) Schematic illustration showing the concept of FBXO22 self-degrader. B) Metabolic conversion of primary diamine to dialdehyde. C) HiBiT-FBXO22 assay results for Jurkat cells treated with the primary diamines for 5 hours. D) Western blots showing FBXO22 degradation in Jurkat cells pre-treated with the indicated inhibitors for 1 hour, followed by treatment with 1 µM hexane-1,6-diamine for 5 hours. E) HiBiT-FBXO22 assay results for Jurkat cells treated with methylated hexane-1,6-diamine for 5 hours. F) Western blots showing FBXO22 degradation in Jurkat cells treated with indicated concentration of AHPC(Me)-C6-NH_2_ for 5 hours or 24 hours. G) Western blots showing protein levels of FBXO22 and its downstream signals in THP-1 cells treated with indicated concentration of hexane-1,6-diamine for 24 hours.

To confirm that hexane-1,6-diamine-induced FBXO22 self-degradation is mediated by UPS, we pretreated the cells with UPS pathway inhibitors as above and found that the FBXO22 degradation was largely blocked. Pretreatment with AG also reduced the FBXO22 degradation, indicating the metabolic conversion to aldehyde analog is required (Figure 4D). Furthermore, methylating one amine group to block its oxidation completely abolished FBXO22 degradation, suggesting that both primary amines must be converted to aldehydes to induce FBXO22 self-degradation (Figure 4E).

Given these results, we next examined whether hexane-1,6-diamine could induce self-degradation of endogenous FBXO22. In Jurkat cells, treatment with hexane-1,6-diamine for 5 hours led to dose-dependent FBXO22 degradation, achieving over 50% reduction at 1 μM. Extending incubation to 24 hours substantially enhanced degradation, resulting in near-complete degradation at the same concentration (Figure 4F). Compared to AHPC(Me)-C6-NH_2_, hexane-1,6-diamine exhibited a more pronounced dependency on incubation time, supporting the idea that metabolic conversion to aldehyde is a rate-determination step, as two deamination steps are required. Moreover, hexane-1,6-diamine did not alter BACH1 protein levels in Jurkat cells but increased BACH1 abundance in THP-1 cells, particularly at 10 μM, where FBXO22 degradation was more robust, indicating that the FBXO22 self-degrader is also biologically active (Figure 4G).

### Charting reactive warheads for hijacking FBXO22

While newly characterized E3 ligases offer opportunities for targeted degradation, their practical utility depends on the availability of effective ligands. Previous studies have documented the use primary alkylamine or alkyl aldehyde as FBXO22 recruiters to degrade targets including NSD2^24^, XIAP^49^, FKBP12^25^ and SMARCA2/A4^28^. FBXO22-mediated BRD4 degradation was successfully achieved using a chloroacetamide-based covalent fragment^26^, however, the primary amine tethering strategy proved ineffective, as confirmed by HiBiT assay results^25^ (Figure S10). Hence, we next sought to explore novel warheads to more effectively harness FBXO22 in TPD applications.

Given that the reaction between thiols and aldehydes to form hemithioacetals is highly reversible, the resulting covalent engagement may lack sufficient stability to support processive ubiquitination. We hypothesized that enhancing the hemithioacetal stability could promote the formation of a FBXO22-containing ternary complex and thereby improve degradation efficacy, as shown in a recent study where a covalent cysteine warhead (chloroacetamide or acrylamide) engages C227/228 to recruit FBXO22 for a broader applicability in TPD^26^. Considering AHPC(Me)-C6-NH_2_ has proven effective for FBXO22 degradation by engaging C326, we installed various warheads on VHL ligand and tested them in the Hibit-FBXO22 assay. We first synthesized acrylamide (ZZ7-24-009) and chloroacetamide (ZZ7-24-010) analogs, two commonly used cysteine targeting covalent warheads. Despite having similar linker geometry to AHPC(Me)-C6-NH_2_, neither compound induced degradation in the Hibit-FBXO22 assay nor formed a ternary complex with the VCB complex and SKP1/FBXO22 (Figure 5A, S11A, B). Given that acrylamides and chloroacetamides are generally more reactive toward cysteine residue than alkyl aldehydes^50^, these results suggested that aldehyde functional group is essential for effective FBXO22 engagement.

**Figure 5.**
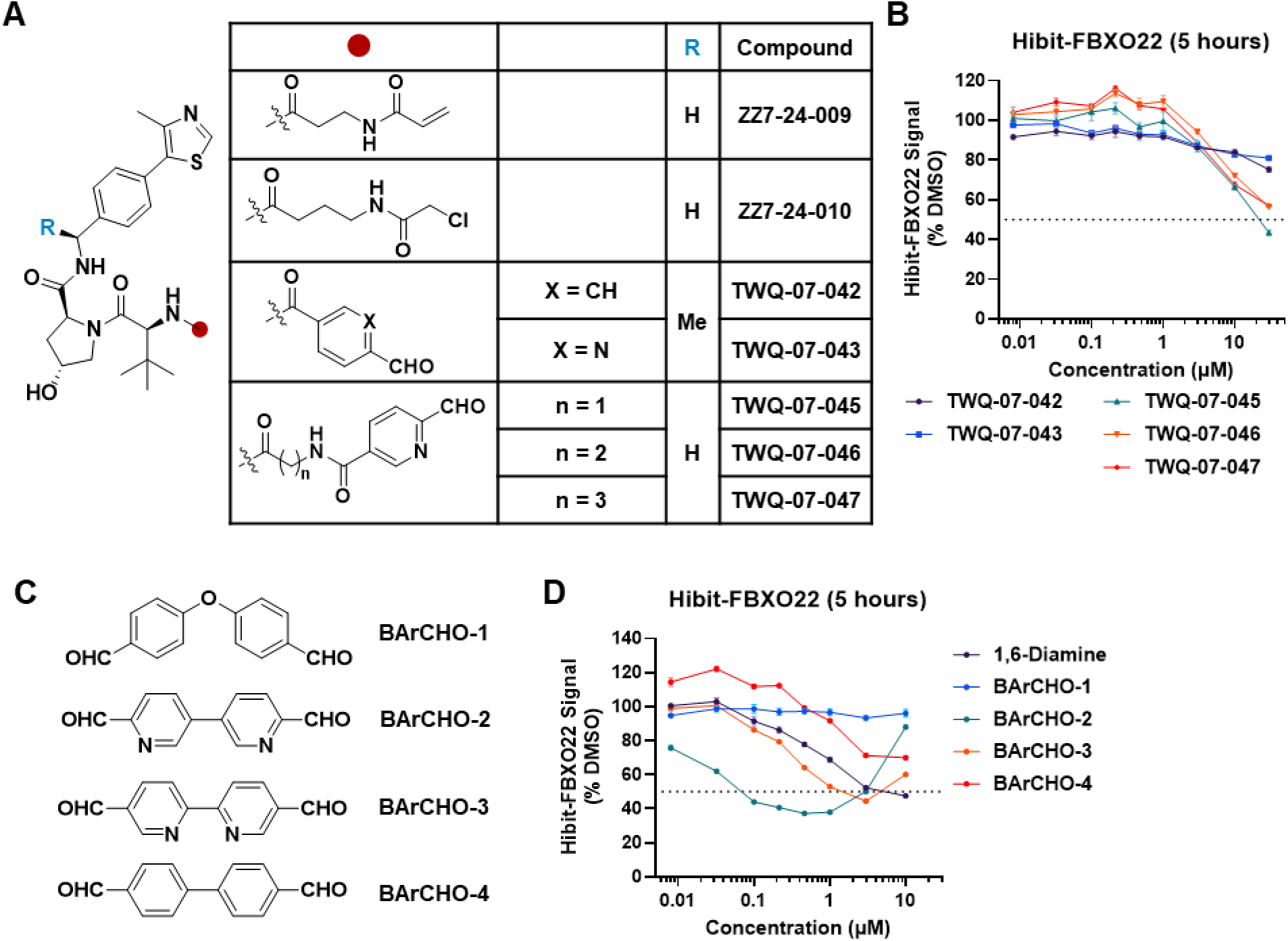
Charting cysteine reactive warhead for targeting FBXO22. A) Chemical structure of VHL ligand tethered with various covalent warheads. B) HiBiT-FBXO22 assay results for Jurkat cells treated with the VHL ligand-based degraders for 5 hours. C) Chemical structure of bi-aromatic dialdehyde (BArCHO). D) HiBiT-FBXO22 assay results for Jurkat cells treated with the BArCHOs for 5 hours.

To broaden the aldehyde repertoire, we focused on aromatic aldehydes, whose reactivity and stability can be better tuned through resonance and electronic effects^51^. We tested two representative aromatic aldehydes: benzaldehyde (BA) and 2-pyridinecarboxaldehyde (2-PCA). In our initial design, we installed aromatic aldehydes on the VHL recruiting ligand. We discovered that compounds with a spacer between VHL ligand and 2-PCA exhibited modest dose-dependent FBXO22 degradation. Among them, TWQ-07-045, which features glycine as spacer, achieved approximately 50% D_max_ in the Hibit-FBXO22 assay (Figure 5A, B, S12A). Encouraged by this, we hypothesized that bi-aromatic dialdehyde (BArCHO) could also facilitate FBXO22 self-degradation (Figure 5C). Indeed, the BArCHO-2 (3,3’-bipyridine-6,6’-dicarbaldehyde) displayed improved self-degradation with a DC_50_ below 0.1 μM and a D_max_ exceeding 60% in the HiBiT-FBXO22 assay. In contrast, bipyridine analogs containing 3-pyridine aldehyde (BArCHO-3) showed lower efficacy, and benzaldehyde derivatives were inactive (Figure 5D, S12B, C). The degradation efficacy correlated with the intrinsic reactivity of the aromatic aldehyde^52^, suggesting that aldehyde reactivity directly influences the FBXO22 self-degradation. Interestingly, despite the high reactivity of 2-PCA, neither BArCHO-2 nor TWQ-07-045 exhibited cytotoxicity (Figure S12D,E). Collectively, these findings identify 2-PCA as a promising alternative reactive warhead for targeting FBXO22.

### 2-pyridinecarboxaldehyde enables FBXO22-mediated targeted protein degradation

2-PCA has been previously reported as a selective N-terminal protein labeling reagent used in protein conjugation chemistry^53^, but its utility in TPD has not been investigated. Building on the promising reactivity of 2-PCA as an FBXO22-directed warhead, we next investigated whether tethering 2-PCA to the BET bromodomain inhibitor JQ1^54^ could enable BRD4 degradation, as JQ1 tethered with a primary alkylamine failed to induce BRD4 degradation^25^. We installed 2-PCA on JQ1 with linkers of varying lengths and measured BRD4 degradation activity using a Hibit-BRD4 assay in Jurkat cells (Figure S13). Remarkably, compounds with alkyl linkers exhibited degradation of BRD4, with ZZ7-17-060 (C3 linker) showing the highest potency, achieving a D_max_ of around 32% (Figure 6A). Treatment with MLN-4924 or MG132 blocked ZZ7-17-060 induced BRD4 degradation, suggesting that the effect is mediated by the UPS. We further synthesized several analogs with the optimal linker for SAR exploration. As expected, a benzaldehyde analog (ZZ7-24-036) showed no activity presumably due to its lower aldehyde reactivity. Similarly, either swapping the amide with a sulfonamide (ZZ7-24-031) or pyridine with pyrimidine (ZZ7-24-030) also abolished the BRD4 degradation activity (Figure S14). We reasoned that these modifications increased the reactivity of the aromatic aldehyde but also reduced its cellular stability, whereas 2-PCA achieves an optimal balance between reactivity and stability.

**Figure 6.**
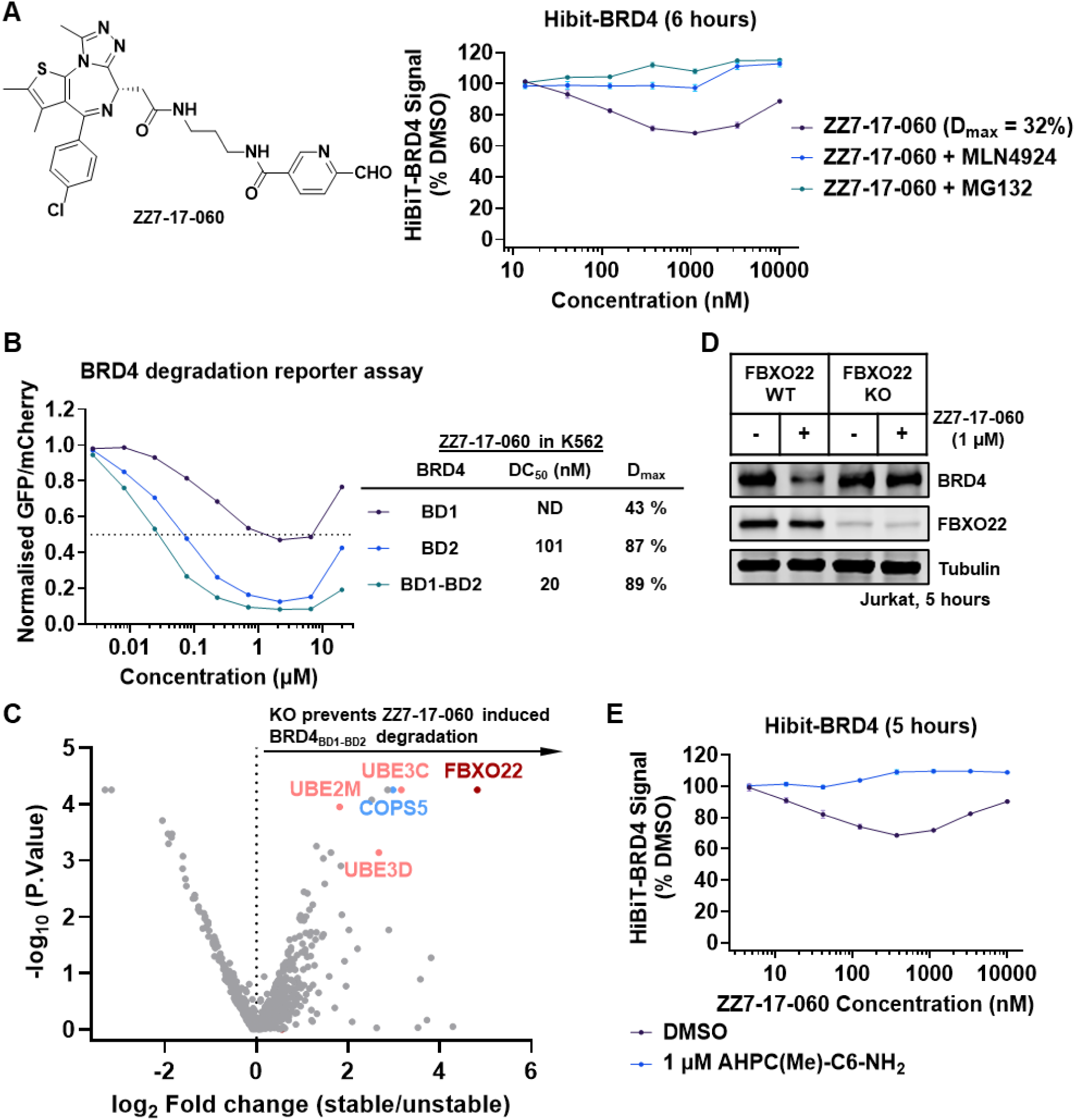
2-pyridinecarboxyaldehyde (2-PCA) enables FBXO22 mediated BRD4 degradation. A) Chemical structure of ZZ7-17-060 and HiBiT-BRD4 assay results for Jurkat cells pre-treated with the indicated inhibitors for 1.5 hours, followed by treatment with ZZ7-17-060 for 6 hours. B) Identifying of the BRD4 region required for ZZ7-17-060 -induced degradation with a cellular fluorescent reporter assay. The examined reporters were either the isolated BRD4 bromodomains (BD1 or BD2) or a tandem construct comprising BD1 and BD2 connected by the intervening native sequence (BRD4_BD1-BD2_). C) Ubiquitin-proteasome system (UPS)-focused CRISPR screen for BRD4_BD1-BD2_-eGFP stability in K562-Cas9 cells treated with 1 μM ZZ7-17-060 for 16 hours. Note the adaptor protein SKP1 is not in the BISON library. D) Western blots showing BRD4 degradation in WT or FBXO22-KO Jurkat cells treated with 1 μM ZZ7-17-060 for 5 hours. E) HiBiT-BRD4 assay results for WT Jurkat cells pre-treated with 1 μM AHPC(Me)-C6-NH_2_, followed by treatment with ZZ7-17-060 for 5 hours.

BRD4 contains two bromodomains (BDs)^55^, and JQ1 binds both BDs with similar affinity^54^. To identify which region of BRD4 is responsible for ZZ7-17-060-induced degradation, we employed a fluorescent reporter assay to monitor BRD4 stability in K-562 cells^56^. Reporters containing either bromodomain 2 (BRD4-BD2) alone or a tandem construct with both BD1 and BD2 showed strong degradation, with D_max_ exceeding 87%. In contrast BRD4-BD1 reporter displayed modest degradation with a D_max_ of approximately 43% (Figure 6B). Thus, ZZ7-17-060 harnesses both BDs, though preferably BRD4-BD2, for BRD4 degradation. Encouraged by the BRD4 degradation efficacy of ZZ7-17-060 in K-562 cells, we next used an unbiased screening approach to assess 2-PCA selectivity for E3 ligase recruitment. To this end, we conducted a fluorescence-activated cell sorting (FACS)–based CRISPR screen, targeting 713 genes involved in the UPS pathway^57^. Remarkably, FBXO22 emerged as the only significantly enriched E3 ligase, along with CUL1 and several candidate E2 enzymes (Figure 6C, S15). Consistent with this, genetic knockout or chemical knockdown of FBXO22 with AHPC(Me)-C6-NH_2_ completely rescued the ZZ7-17-060-induced BRD4 degradation (Figure 6D, E, S16).

To further elucidate the mechanism of ZZ7-17-060, and more broadly to assess whether 2-PCA directly binds to FBXO22, we investigated their interaction in more detail. First, we tested whether simple 2-PCA derivatives, 2-formylpyridine and PCA-alkyne, could compete the activity of ZZ7-16-060 in inducing BRD4 degradation. Pretreatment with either compound completely abolished ZZ7-16-060 -induced BRD4 degradation, suggesting that 2-PCA is capable of directly engaging FBXO22 (Figure 7A). Next, to confirm the formation of a new protein-protein interaction, we incubated ZZ7-17-060 with HEK293T lysates expressing FLAG-tagged BRD4 and observed enrichment of FBXO22 only in the treatment group (Figure 7B). Further, we conducted similar IP experiments to test whether 2-PCA engages the same cysteine as the primary aldehyde. We found that the C326A mutation completely prohibited the FBXO22 engagement (Figure 7C). Interestingly, we noticed Y390 positioned above C326, which may act as a lid over the putative binding pocket. Indeed, the Y390A mutation also completely blocked the FBXO22 engagement, suggesting that this tyrosine may contribute π-π interactions with the pyridine ring of 2-PCA, thereby facilitating its binding to FBXO22 (Figure 7C,D).

**Figure 7.**
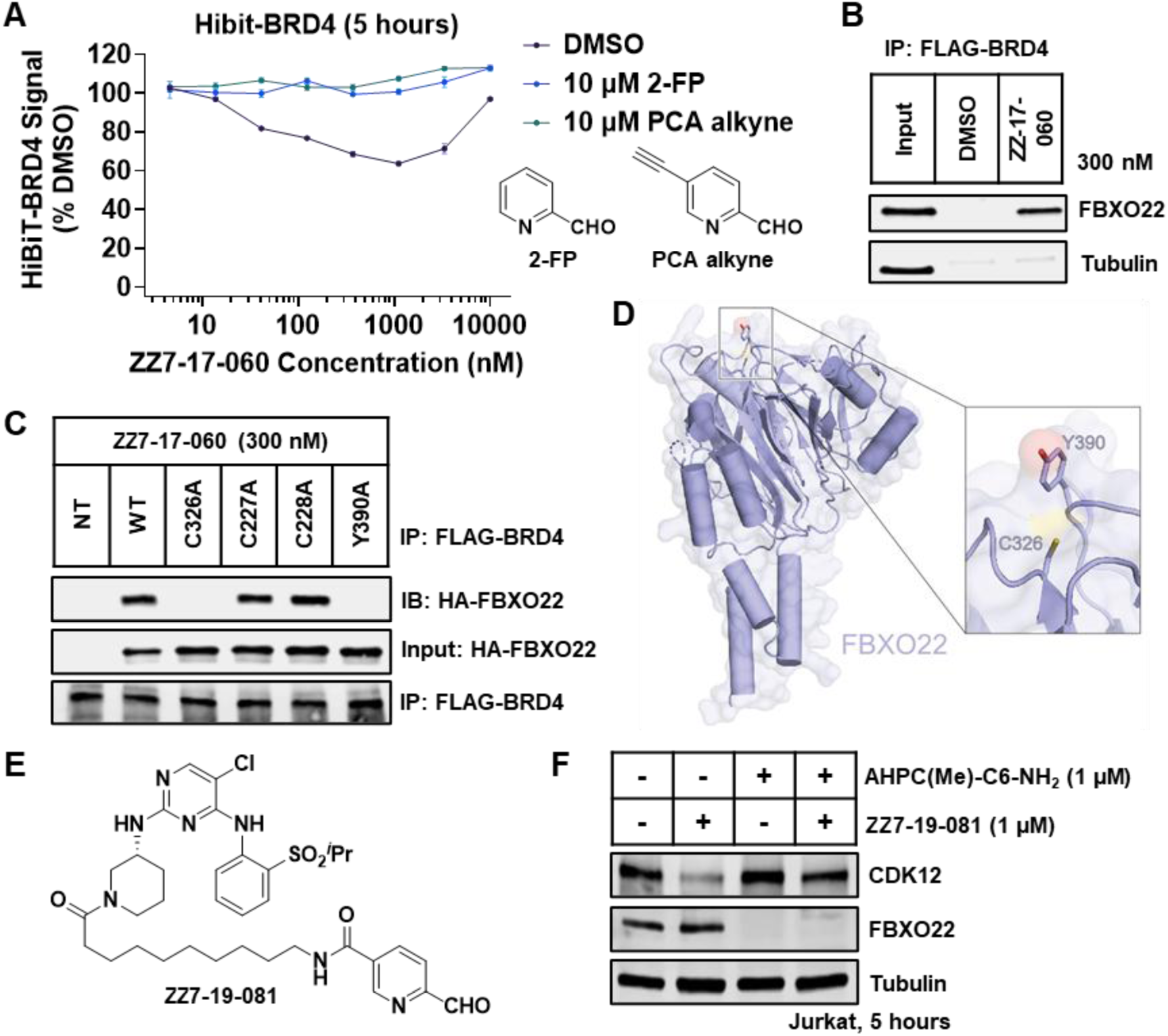
2-PCA engages FBXO22 Cys326 to degrade target protein. A) HiBiT-BRD4 assay results for Jurkat cells pre-treated with 10 μM 2-FP or PCA-Alkyne for 1 hour, followed by treatment with ZZ7-17-060 for 5 hours. B) Co-immunoprecipitation of FLAG-tagged BRD4 and endogenous FBXO22 in the presence of 300 nM ZZ7-17-060. C) Co-immunoprecipitation of FLAG-tagged BRD4 and HA-FBXO22 (WT or mutations) in the presence of 300 nM ZZ7-17-060. NT indicates no transfection of FBXO22 constructs. D) Schematic illustration of 2-PCA putative binding pocket on FBXO22 structure (PDB:8S7E). Cys 326 and Tyr 390 are highlighted. E) Chemical structure of CDK12 degrader ZZ7-19-081. E) Western blots showing CDK12 degradation in Jurkat cells pre-treated with 1 μM AHPC(Me)-C6-NH_2_ for 1 hour, followed by treatment with 1 µM ZZ7-19-081 for 5 hours.

Finally, to probe the generality of FBXO22 recruitment to diverse targets, we tethered 2-PCA to a parental CDK12 inhibitor from the selective CDK12 degrader BSJ-4-116 via an alkyl linker (Figure 7E)^58^. The resulting compound, ZZ7-19-081, exhibited modest CDK12 degradation, which was rescued by chemical knockdown of FBXO22 with AHPC(Me)-C6-NH_2_, indicating that FBXO22 mediates ZZ7-19-081-induced CDK12 degradation (Figure 7F). Together, these results further support 2-PCA as a selective degradation warhead that recruits FBXO22 for TPD.

## Discussion

Recent reports investigating the mode of action of serendipitously discovered degraders have shown that FBXO22 can be hijacked for TPD using small molecules that function as prodrugs - compounds metabolically converted in cells to reactive aldehydes, enabling covalent engagement with FBXO22^24-28^. These studies established proof of concept for FBXO22 as a ligandable E3 ligase but left open critical questions regarding direct engagement strategies, degradation selectivity, and the broader applicability of FBXO22 in induced proximity approaches.

In this work, we expand and deepen this emerging paradigm by developing the first direct chemical degraders of FBXO22 and systematically exploring their mechanisms of action. Our lead compound, AHPC(Me)-C6-NH_2_, is a potent and selective degrader that mimics genetic loss of FBXO22 and serves as a chemical biology tool for probing its function. Comprehensive mechanistic studies—including SAR exploration, pathway inhibitor rescue experiments, and ternary complex formation assays—revealed its activity requires metabolic conversion of the primary amine into an alkyl aldehyde, consistent with prior reports that aldehyde-mediated covalent engagement enables FBXO22 recruitment. Despite its structural simplicity, AHPC(Me)- C6-NH_2_ exhibits remarkable proteome-wide selectivity for FBXO22 degradation. We hypothesized that this selectivity stems partially from the preferential protein-protein interaction between FBXO22 and the VHL upon ternary complex formation, as FBXO22 emerges as the predominant protein in proximity to VHL upon treatment with the AHPC-CHO. In addition, the aldehyde engages Cys326 of FBXO22 in a reversible covalent manner, whereas stronger covalent warheads failed to induce ternary complex formation or FBXO22 degradation, suggesting that FBXO22 selectivity also hinges on a finely balanced reversible engagement.

Building on this, we explored the concept of homo-PROTAC-mediated self-degradation, wherein E3 ligases are dimerized to induce their own ubiquitination and proteasomal degradation, and identified hexane-1,6-diamine as a minimal self-degrader of FBXO22. In contrast, biologically relevant shorter-chain diamines^59, 60^ such as putrescine (1,4-diaminobutane) and cadaverine (1,5-diaminopentane) lacked this activity. The inability of these endogenous short-chain diamines to trigger FBXO22 degradation matches with their benign cellular roles and emphasizes the necessity of appropriate chain length and the related alkyl dialdehyde reactivity for effective FBXO22 recruitment and self-degradation. These insights mirror prior homo-PROTAC strategies used for VHL^23^, CRBN^48^, and MDM2^61^ and highlight a potentially generalizable approach for chemically modulating E3 ligase levels. Given FBXO22’s frequent overexpression in diverse cancers, the selective degraders we describe may offer starting points for therapeutic targeting of FBXO22-driven tumorigenic processes.

A key advance of our study is the discovery of 2-pyridinecarboxaldehyde (2-PCA) as a metabolically-independent reactive warhead for FBXO22 recruitment. Unlike primary alkylamines, 2-PCA engages FBXO22 directly without requiring metabolic activation, enabling the efficient FBXO22 self-degradation and degradation of diverse targets such as BRD4 and CDK12. This not only broadens the arsenal of electrophilic warheads available for TPD but also highlights the potential of aromatic aldehydes in designing selective, reversible E3 ligase recruiters. The effectiveness of 2-PCA–based degraders was reinforced by unbiased CRISPR screening, which confirmed FBXO22 as the sole E3 ligase required for degradation of the model protein BRD4. Given FBXO22’s frequent overexpression in tumors, leveraging it as a recruiter may bias degrader activity toward cancer cells, offering a potential therapeutic advantage.

These findings collectively position FBXO22 as both a degradable protein target and a promising recruiter for induced proximity approaches. The toolkit we provide - comprising selective FBXO22 degraders, homo-PROTACs, and aromatic aldehyde-based warheads - complements and extends prior prodrug-based strategies by enabling direct, modular chemical control of FBXO22. Our results also highlight the broader principle that reversible covalent engagement, guided by intrinsic E3 ligase reactivity and proximity preferences, can unlock new opportunities for E3 ligase harnessing in TPD.

Despite 2-PCA and its derivatives having been reported to react specifically with N-terminal amino acid in peptides or proteins^62, 63^ at high concentrations (∼ 5 mM), their utility in cysteine targeting remains rarely documented^64^. Our mutation analysis suggests that 2-PCA–based degraders recruit FBXO22 through Cys326 at substantially lower concentrations. Supporting this, both 2-FP and PCA-Alkyne fully rescued BRD4 degradation, suggesting that 2-PCA may directly bind to FBXO22. Attempts to capture the binding between FBXO22 and 2-PCA were not successful, likely due to the reversibility of hemithioacetal linkage. Nevertheless, the robust neo-protein–protein interaction induced by ZZ7-17-060 and its potent activity against BRD4-BD2 support the feasibility of determining the structure of BRD4-BD2/ZZ7-17-060/FBXO22 complex.

Looking forward, structural elucidation of FBXO22–ligand complexes and medicinal chemistry optimization of aromatic aldehyde warheads will be key to enhancing selectivity, potency, and substrate scope. Such advances could ultimately establish FBXO22 as a versatile alternative to canonical recruiter ligases like CRBN and VHL, expanding the reach of TPD approaches in both mechanistic studies and therapeutic applications.

## Methods

### General cell biology methods

Roswell Park Memorial Institute (RPMI) 1640 medium and Dulbecco’s modified Eagle’s medium (DMEM), Iscove’s Modified Dulbecco’s Medium (IMDM), Heat-inactivated Fetal bovine serum (FBS), penicillin-streptomycin (10,000 units/mL sodium penicillin G and 10,000 μg/mL streptomycin), trypsin-EDTA solution (1×), and phosphate-buffered saline (PBS; 1×) were purchased from Gibco Invitrogen Corp (Grand Island, NY, USA). Eagle’s Minimum Essential Medium (EMEM) and DMSO were purchased from ATCC (Manassas, Virginia, USA). MG132, MLN-4924, Bortezomib, and dBET6 were purchased from MedChemExpress (Monmouth Junction, NJ, USA). All other chemicals were purchased from Sigma-Aldrich (St. Louis, MO, USA), unless indicated otherwise.

### Cell lines

HEK293T, K562, Kelly, MOLT-4, HepG2, Jurkat and THP-1 cells were purchased and verified from the ATCC. HEK293T cells were cultured in DMEM medium, with 10% heat-inactivated FBS and 1% penicillin-streptomycin. Kelly, MOLT-4, Jurkat, K562 and THP-1 cells were cultured in RPMI medium, with 10% FBS and 1% penicillin-streptomycin. HepG2 cells were cultured in EMEM medium, with 10% FBS, 1% penicillin-streptomycin. All cell lines were grown in humidified tissue culture incubators at 37 ℃ with 5% CO_2_ and were mycoplasma-negative based on monthly testing using the MycoAlert mycoplasma detection kit (Lonza, Basel, Switzerland). For all experiments, cells had undergone fewer than 18 passages.

### Western blotting analysis

The cells were washed once with DPBS before lysing in RIPA lysis buffer (50 mM Tris, 150 mM NaCl, 0.5% deoxycholate, 0.1% SDS, 1.0% Triton X-100, pH 7.4) (BP-116TX, Boston Bioproducts, Inc., Milford, MA, USA) supplemented with a protease inhibitor cocktail (11836153001, Roche) for 15 min on ice. The lysates were centrifuged at 13,000g (4 °C, 20 minutes), and the supernatant was collected. Protein concentrations of the cell lysates were quantified using the BCA method and a BCA Protein Assay Kit (Thermo Fisher Scientific, Waltham, MA, USA). Equal amounts of protein were mixed with 6x Laemelli sample buffer (J61337.AC, Thermo Fisher Scientific) containing 150 mM DTT and then subjected to 4-20% sodium dodecyl sulfate polyacrylamide gel electrophoresis (SDS-PAGE) and transferred to nitrocellulose membranes (cat. no. 1620112; Bio-Rad Laboratories, Hercules, CA, USA). The membranes were blocked using Intercept® (TBS) Blocking Buffer (LI-COR Biosciences, Lincoln, NE, USA) and subsequently probed with appropriate primary antibodies [anti-BRD4 (cat. no. ab128874; Abcam, Cambridge, UK and cat. no. 63759; Cell Signaling Technology, Danvers, MA, USA), anti-α-Tubulin (cat. no. 3873; Cell Signaling Technology), anti-FLAG M2 (cat. no. F1804; Sigma-Aldrich), anti-FBXO22 (13606-1-AP, Proteintech, Rosemont, IL, USA), anti-CRBN (71810S, Cell Signaling Technology, Danvers, MA, USA), anti-VHL (68547S, Cell Signaling Technology, Danvers, MA, USA), anti-BACH1 (F-9) (sc-271211, Santa Cruz Biotechnology, Inc., Dallas, Texas, USA), anti-CDK12 (VMA00874, Bio-Rad Laboratories, Hercules, CA, USA)] at 4 °C overnight. The membranes were washed with TBST (20 mM Tris, pH 7.5, 150 mM NaCl, 0.1% Tween-20) (IBB-180X, Boston Bioproducts, Inc., Milford, MA, USA) for 4*5 min at room temperature and incubated with IRDye 800-labeled goat anti-rabbit IgG (LI-COR Biosciences, cat. no. 926-32211) or IRDye 680RD goat anti-Mouse IgG (LI-COR Biosciences, cat. no. 926-68070) secondary antibodies at room temperature for 1 h. After washing the membranes with TBST for 4*5 min at room temperature, the membranes were detected on Li-COR Odyssey CLx system.

### Generation of Hibit-FBXO22 stable transfected Jurkat cells

FBXO22 (Uniprot: Q8NEZ5-1) was fused with the HiBiT-RR sequence (VSGWRLFRRIS)^45^ linked to the protein of interest via a Gly-Ser linker. Codon-optimized coding sequences were ordered as gBlock fragments (IDT) and cloned into vector pN103 under the control of the EF1a promoter. Corresponding lentivirus was used to transduce Jurkat cells cultured as described above, and transduced cells were selected by growth in 2 µg/ml puromycin added directly to the culture medium. Pooled selectants were verified for kinase expression using the Nano-Glo HiBiT Lytic Detection System (Promega) as described by the manufacturer.

### Hibit-FBXO22 assay

HiBiT-FBXO22 Jurkat cells were seeded into 384-well white plates (5,000 cells/well for 5h treatment; 7,500 cells/well for 18h treatment) and treated with compounds in triplicate. After given time of treatment, the plates were subjected to Nano-Glo HiBiT Lytic Detection System as described in manufacturer’s manual. IC_50_ values were determined using a nonlinear regression curve fit in GraphPad PRISM v10.5.0. In the pathway inhibitor rescue experiments, cells were pretreated for 1 hour before the 5-hour incubation with test compound. In the media switching experiments, cells were seeded in Opti-MEM (Gibco) and treated with compounds. For 18h treatment experiments with BArCHO compounds, the HiBiT signal was normalized to the Cell Titer Glo cell viability signal for each matching condition and to DMSO for each treatment. For other experiments, the HiBiT signal was normalized to DMSO for each treatment.

### Whole cell quantitative proteomics cell treatments

HEK293T and Kelly cells were treated with 1 µM and 5 µM ZXH-3-122, respectively, or DMSO vehicle control for 6 hours and 10 hours, respectively.

### Whole cell quantitative proteomics sample processing

Cells were lysed by the addition of lysis buffer (8 M urea, 50 mM NaCl, 50 mM 4-(2-hydroxyethyl)-1-piperazineethanesulfonic acid (EPPS) pH 8.5, Protease and Phosphatase inhibitors) followed by manual homogenization by 20 passes through a 21-gauge (1.25 in. long) needle or homogenization by bead beating (BioSpec) for three repeats of 30 seconds at 2400 strokes/min. Lysate was clarified by centrifugation and protein quantified using Bradford (Bio-Rad) assay. 50-100 µg of protein for each sample was reduced, alkylated and precipitated using methanol/chloroform as previously described^65^ and the resulting washed precipitated protein was allowed to air dry. Precipitated protein was resuspended in 4 M urea, 50 mM HEPES pH 7.4, buffer for solubilization, followed by dilution to 1 M urea with the addition of 200 mM EPPS, pH 8. Proteins were digested for 12 hours at room temperature with LysC (1:50 ratio), followed by dilution to 0.5 M urea and a second digestion step was performed by addition of trypsin (1:50 ratio) for 6 hours at 37 °C or digested with the addition of LysC (1:50; enzyme:protein) and trypsin (1:50; enzyme:protein) for 12 h at 37 °C.

### TMT quantitative proteomics and data analysis

Anhydrous ACN was added to each peptide sample to a final concentration of 30%, followed by addition of 10-plex Tandem mass tag (TMT) reagents at a labelling ratio of 1:4 peptide:TMT label. TMT labelling occurred over a 1.5 hour incubation at room temperature followed by quenching with the addition of hydroxylamine to a final concentration of 0.3%. Each of the samples were combined using adjusted volumes and dried down in a speed vacuum followed by desalting with C18 SPE (Sep-Pak, Waters). The sample was offline fractionated into 96 fractions by high pH reverse-phase HPLC (Agilent LC1260) through an aeris peptide xb-c18 column (phenomenex) with mobile phase A containing 5% acetonitrile and 10 mM NH_4_HCO_3_ in LC-MS grade H_2_O, and mobile phase B containing 90% acetonitrile and 5 mM NH_4_HCO_3_ in LC-MS grade H_2_O (both pH 8.0). The resulting 96 fractions were recombined in a non-contiguous manner into 24 fractions and desalted using C18 solid phase extraction plates (SOLA, Thermo Fisher Scientific) followed by subsequent mass spectrometry analysis.

Data were collected using an Orbitrap Fusion Lumos mass spectrometer (Thermo Fisher Scientific, San Jose, CA, USA) coupled with a Proxeon EASY-nLC 1200 LC system (Thermo Fisher Scientific, San Jose, CA, USA). Peptides were separated on a 50 cm 75 μm inner diameter EasySpray ES903 microcapillary column (Thermo Fisher Scientific). Peptides were separated over a 190 min gradient of 6 - 27% acetonitrile in 1.0% formic acid with a flow rate of 300 nL/min.

Qualification was performed using an MS3-based TMT method as described previously^66^. The data were acquired using a mass range of m/z 340 – 1350, resolution 120,000, AGC target 5 x 10^5^, maximum injection time 100 ms, dynamic exclusion of 120 seconds for the peptide measurements in the Orbitrap. Data dependent MS2 spectra were acquired in the ion trap with a normalized collision energy (NCE) set at 35%, AGC target set to 1.8 x 10^4^ and a maximum injection time of 120 ms. MS3 scans were acquired in the Orbitrap with HCD collision energy set to 55%, AGC target set to 2 x 10^5^, maximum injection time of 150 ms, resolution at 50,000 and with a maximum synchronous precursor selection (SPS) precursors set to 10.

Proteome Discoverer 2.2 (Thermo Fisher Scientific) was used for .RAW file processing and controlling peptide and protein level false discovery rates, assembling proteins from peptides, and protein quantification from peptides. The MS/MS spectra were searched against a Swissprot human database (January 2017) containing both the forward and reverse sequences. Searches were performed using a 10-ppm precursor mass tolerance, 0.6 Da fragment ion mass tolerance, tryptic peptides containing a maximum of two missed cleavages, static alkylation of cysteine (57.0215 Da), static TMT labelling of lysine residues and N-termini of peptides (229.1629 Da), and variable oxidation of methionine (15.9949 Da). TMT reporter ion intensities were measured using a 0.003 Da window around the theoretical m/z for each reporter ion in the MS3 scan. The peptide spectral matches with poor quality MS3 spectra were excluded from quantitation (summed signal-to-noise across channels < 100 and precursor isolation specificity < 0.5), and the resulting data was filtered to only include proteins with a minimum of 2 unique peptides quantified. Reporter ion intensities were normalized and scaled using in-house scripts in the R framework (R: A language and environment for statistical computing (R Foundation for Statistical Computing, 2014).). Statistical analysis was carried out using the limma package within the R framework^67^.

### IP-MS

A total of 1 × 10^7^ cells per IP were collected and lysed in lysis buffer (50 mM Tris pH 8, 200 mM NaCl, 2 mM TCEP, 0.1% NP-40, 10 units turbonuclease/200 µL buffer, 1x cOmplete protease inhibitor tablet/5 mL buffer) and sonicated on ice for 5 rounds of 2 seconds followed by 10 second pauses at 25% amplitude. After centrifugation clarification, lysate was transferred to new lobind tubes. 20 µg of Flag-VHL-EloB/C, 1 µM of MLN4924 and CSN5i-3 (neddylation trap)^68^ and 1 µM of selected degraders or DMSO vehicle control were added to each lysate and incubated with end-over-end rotation for 1 hour in the cold room. 20 µL of pre-washed and resuspended M2-Flag magnetic bead slurry was added to each IP and incubated with end-over-end rotation for 1 hour in the cold room. Beads were washed and processed following Baek et al^47^. In brief, beads were washed three times with detergent-containing buffer followed by three washes with non-detergent buffer. Samples were then eluted with 0.1 M Glycine-HCl, pH 2.7. Tris (1M, pH 8.5), followed by reduction, alkylation, and trypsin digestion. Sample digests were acidified with formic acid to a pH of 2-3 prior to desalting using C18 solid phase extraction plates (SOLA, Thermo Fisher Scientific). Desalted peptides were dried in a vacuum-centrifuged and reconstituted in 0.1% formic acid for LC-MS analysis.

### Label free quantitative proteomics for whole cell and IP-MS

Sample digests were acidified with formic acid to a pH of 2-3 before desalting using C18 solid phase extraction plates (SOLA, Thermo Fisher Scientific). Desalted peptides were dried in a vacuum-centrifuged and reconstituted in 0.1% formic acid for liquid chromatography-mass spectrometry analysis. Data were collected using a TimsTOF HT (Bruker Daltonics, Bremen, Germany) coupled to a nanoElute2 LC pump (Bruker Daltonics, Bremen, Germany) via a CaptiveSpray nano-electrospray source. Peptides were separated on a reversed-phase C_18_ column (25 cm x 75 µm ID, 1.6 µM, IonOpticks, Australia) containing an integrated captive spray emitter. Peptides were separated using a 50 min gradient of 2 - 30% buffer B (acetonitrile in 0.1% formic acid) with a flow rate of 250 nL/min and column temperature maintained at 50 °C.

The TIMS elution voltages were calibrated linearly with three points (Agilent ESI-L Tuning Mix Ions; 622, 922, 1,222 *m/z*) to determine the reduced ion mobility coefficients (1/K_0_). To perform diaPASEF, we used py_diAID^69^, a python package, to assess the precursor distribution in the *m/z*-ion mobility plane to generate a diaPASEF acquisition scheme with variable window isolation widths that are aligned to the precursor density in m/z. Data was acquired using twenty cycles with three mobility window scans each (creating 60 windows) covering the diagonal scan line for doubly and triply charged precursors, with singly charged precursors able to be excluded by their position in the m/z-ion mobility plane. These precursor isolation windows were defined between 350 - 1250 *m/z* and 1/k0 of 0.6 - 1.45 V.s/cm^2^.

The diaPASEF raw file processing and controlling peptide and protein level false discovery rates, assembling proteins from peptides, and protein quantification from peptides were performed using library free analysis in DIA-NN 1.8^70^ searched against a Swissprot human database (January 2021). Database search criteria largely followed the default settings for directDIA including tryptic with two missed cleavages, carbamidomethylation of cysteine, and oxidation of methionine and precursor Q-value (FDR) cut-off of 0.01. Precursor quantification strategy was set to Robust LC (high accuracy) with RT-dependent cross run normalization.

For whole cell samples, proteins with low sum of abundance (<2,000 x no. of treatments) were excluded from further analysis and resulting data was filtered to only include proteins that had a minimum of 3 counts in at least 4 replicates of each independent comparison of treatment sample to the DMSO control.

For IP-MS samples, resulting data was filtered to only include proteins that had a minimum of 3 counts in at least 4 replicates of each independent comparison of treatment sample to the DMSO control. Protein abundances were globally normalized using in-house scripts in the R framework (R Development Core Team, 2014).

For both sample types, proteins with missing values were imputed by random selection from a Gaussian distribution either with a mean of the non-missing values for that treatment group or with a mean equal to the median of the background (in cases when all values for a treatment group are missing)^47^. Protein abundances were scaled and significant changes comparing the relative protein abundance of these treatments to DMSO control comparisons were assessed by two-sided moderated t-test as implemented in the limma package within the R framework^67^.

### Cloning and Protein Expression

His-TEV-VHL (residues 53-C), ELOB (residues N-104), and ELOC (residues 17-C) were co-expressed in LOBSTR BL21(DE3) *E. coli*, purified by His-affinity, followed by liberation of the His-tag by overnight TEV protease incubation at 4C, ion exchange chromatography, and size exclusion chromatography into buffer 25mM HEPES pH 7.5, 200mM NaCl, and 1mM TCEP, pH 7.5. Baculovirus was generated in *Spodoptera frugiperda* cells for SKP1-FBXO22 using pAC8 Strep-Avi-TEV-FBXO22 and pLIB SKP1, and protein was expressed in *Trichoplusia ni* cells by Strep-affinity, followed by ion exchange chromatography and size exclusion chromatography as above.

### Time-Resolved Fluorescence Resonance Energy Transfer (TR-FRET)

VHL-ELOB/C was labeled using BODIPY-NHS at room temperature for 1hr, quenched with 100mM Tris pH 8.0, and desalted into buffer 25mM HEPES pH 7.5, 200mM NaCl, and 1mM TCEP, pH 7.5. Strep-Avi-TEV-FBXO22-SKP1 was biotinylated with BirA, 50mM Biotin, and 5mM ATP, 5mM MgCl2. Ternary complex formation of BODIPY-labeled VHL-ELOBC, SKP1-FBXO22, and compounds were measured by TR-FRET. 50 nM biotinylated SKP1-FBXO22, 2 nM Tb (CoraFluor-1)-labeled Streptavidin (R&D Systems, Cat#7920/20U), and 100 nM BODIPY-VHL-ELOBC were added in assay buffer (25 mM HEPES, 100 mM NaCl, 0.5% Tween-20 (Sigma-Aldrich, Cat#9005-64-5), and 0.5% BSA (Cell Signaling, Cat#9998S)). The reaction mix was added to a 384 well microplate with a final volume of 15 µL, and compounds were titrated to indicated concentrations using a D300e Digital Dispenser (HP). Once compound was added and mixed well, the plate was incubated at room temperature for 1 hr and was measured using a PHERAstar FS microplate reader (BMG). TR-FRET ratio of 520 nm/490 nm was measured as a readout of ternary complex formation, and each datapoints were plotted in Graphpad Prism (v10.1.1).

### Generation of FBXO22 knockout cells

sgRNA constructs were assembled according to standard procedures as described for BsmBI-cut pXPR_023 (gift from S. Corsello; see Addgene #202447 for example). The destination lentiviral transfer vector contains an sgRNA cloning site (U6 promoter) and Cas9-FLAG-P2A-PuroR (EF1alpha promoter). FBXO22 sgRNA sequences are provided below.

>FBXO22_sg_f

CACCGCGCCAGGTTACTCAACACGA

>FBXO22_sg_r

AAACTCGTGTTGAGTAACCTGGCGC

HEK293T cells were transfected with pXPR_023/FBXO22, along with pMD2.G (Addgene #12259) and psPAX2 (Addgene #12260), using Lipofectamine 2000 (Invitrogen) to produce lentivirus. After 48 hours of transfection, the supernatant was collected, filtered, and added to WT Jurkat or HiBiT-BRD4 Jurkat cells in the presence of 5 µg/mL polybrene (APExBIO, cat. no. K2701). Following 48 hours of infection, the infected cells were passaged and cultured in medium containing 1 µg/mL puromycin (Gibco, cat. no. A11138-03) for three days. The surviving cells were grown for an additional five days. The knockout was confirmed by immunoblotting using anti-FBXO22 antibody.

### Hibit-BRD4 assay

The generation of HiBiT-BRD4 Jurkat cells was reported previously^71^. Endogenous BRD4 protein levels were evaluated using the Nano-Glo HiBiT Lytic Detection System (Promega). In brief, 1.5 × 10^4^ HiBiT-BRD4 Jurkat cells were seeded into 384-well plates and incubated with the indicated concentrations of compounds. After 5 h, the plates were subjected to Nano-Glo HiBiT Lytic Detection System as described in manufacturer’s manual. The HiBiTBRD4 assays were conducted in biological triplicates. IC50 values were determined using a nonlinear regression curve fit in GraphPad PRISM v10.3.1.

### BD domain flow cytometry analysis

K562-Cas9 cells expressing the Brd4(BD1)_eGFP_, Brd4(BD2)_eGFP_ or Brd4(BD1-BD2)_eGFP_ degradation reporters were resuspended at 0.7 × 10^6^ ml^−1^ and 50 µl of cell suspension was seeded in 384-well plates. Shortly after, cells were treated with DMSO (n=2) or drug (n=2) for 16 h. The indicated drugs were dispensed with a D300 digital dispenser (Tecan). The fluorescent signal was quantified by flow cytometry (FACSymphony flow cytometer, BD Biosciences). Using FlowJo (flow cytometry analysis software, BD Biosciences), the geometric mean of the eGFP and mCherry fluorescent signal for round and mCherry-positive cells was calculated. The ratio of eGFP to mCherry was normalized to the average of two DMSO-treated controls. Dose-response curves were generated using Prism 10 (GraphPad), and IC50 values were calculated accordingly.

### Bison CRISPR screen for BRD4 stability

The Bison CRISPR library targets 713 E1, E2 and E3 ubiquitin ligases, deubiquitinases and control genes and contains 2,852 guide RNAs (Addgene #169942)^72^. 10% (v/v) of the Bison CRISPR library was added to 10 × 10^6^ BRD4(BD2)_eGFP_ or BRD4(BD1-BD2)_eGFP_ K562-Cas9 cells and transduced (2,400 rpm, 2 h, 37 °C). Eight days later, cells were treated with drug or DMSO for 16 hours and four populations were collected (top 5%, top 5–10%, lowest 5–10% and lowest 5%) based on the BRD4_eGFP_ to mCherry mean fluorescent intensity (MFI) ratio on an MA900 Cell Sorter (Sony, Minato City, Japan). Sorted cells were collected by centrifugation, subjected to direct lysis, and amplified as described before^57^. Amplified sgRNAs were quantified using the Illumina NovaSeq SP platform (Genomics Platform, Broad Institute). The screen was analyzed by comparing stable populations (top 5% eGFP/mCherry expression) to unstable populations (lowest 5% eGFP/mCherry expression) as described before^57^.

### Pull-down assay

HEK293T cells were transfected with pcDNA5-Flag-BRD4-WT (Addgene #90331) using Lipofectamine 2000. After 36h, cell lysates were prepared in NP-40 IP lysis buffer (50 mM Tris, 150 mM NaCl, 5 mM EDTA, 1% Nonidet P-40, pH 7.4) (BP-119, Boston Bioproducts, Inc., Milford, MA, USA) supplemented with a protease inhibitor cocktail and then centrifuged at 20,000g for 15 min at 4 °C. The lysate’s concentration was then measured using the BCA assay (ThermoFisher), and small amount of lysate was saved for the “Input”. 200 μg of lysates were incubated with either tested compounds or DMSO as a control at room temperature for 1h. Following treatment, 10 μL Anti-FLAG® M2 magnetic beads (Millipore-sigma) were added to the lysates and incubated overnight at 4 °C with gentle rotation. The beads were washed once with NP-40 IP lysis buffer and twice with 0.02% Tween-20 in PBS to remove nonspecific proteins. The enriched protein was eluted with 30 μL of 50 mM glycine (pH 2.8) for 5 min at room temperature and brought to neutral with 2 μL of 1 M Tris-base, “output”. Both input and output protein were mixed with 6x Laemelli sample buffer (Alfa Aesar) containing 150 mM DTT, then analyzed by SDS-PAGE and Western Blot.

### Compound resources

Compound resources are provided in the Supplementary Methods (Chemical compounds)

### Chemical synthesis

Synthesis details are provided in the Supplementary Methods (Chemical synthesis).

## Data availability

Data generated in this study are provided in the manuscript, supplementary information, and source data files. Additional data supporting the findings of this study are available from the corresponding author upon reasonable request.

## Acknowledgements

This work was support by the National Institutes of Health (NIH) grants R01CA218278 (to N.S.G. and E.S.F.), R01CA262188 and R01CA214608 (both E.S.F.), NIH High End Instrumentation grant (1S10OD028697-01) (to N.S.G.), departmental funds from Stanford Chemical and Systems Biology and Stanford Cancer Institute (to N.S.G.), and Basic Science Research Program through the National Research Foundation of Korea (NRF) funded by the Ministry of Education (RS-2024-00410290) (to W.S.B.). B.L.E. received support for this work from the NIH (P01CA066996) and the Howard Hughes Medical Institute. Z.K. was supported by a Swiss National Science Foundation Postdoc.Mobility grant (P500PB_214385). K.B. is a Meghan E. Raveis Fellow of the Damon Runyon Cancer Research Foundation (DRG-2514-24). T.Q. thanks Dr. Stephen M. Hinshaw for helpful discussions.

