## Supplementary Figures for "Development of Degraders and 2-pyridinecarboxyaldehyde (2-PCA) as a recruitment Ligand for FBXO22"

#### SI-1. Chemo-proteomics reveal unexpected degradation of FBXO22 by a CRBN based degrader

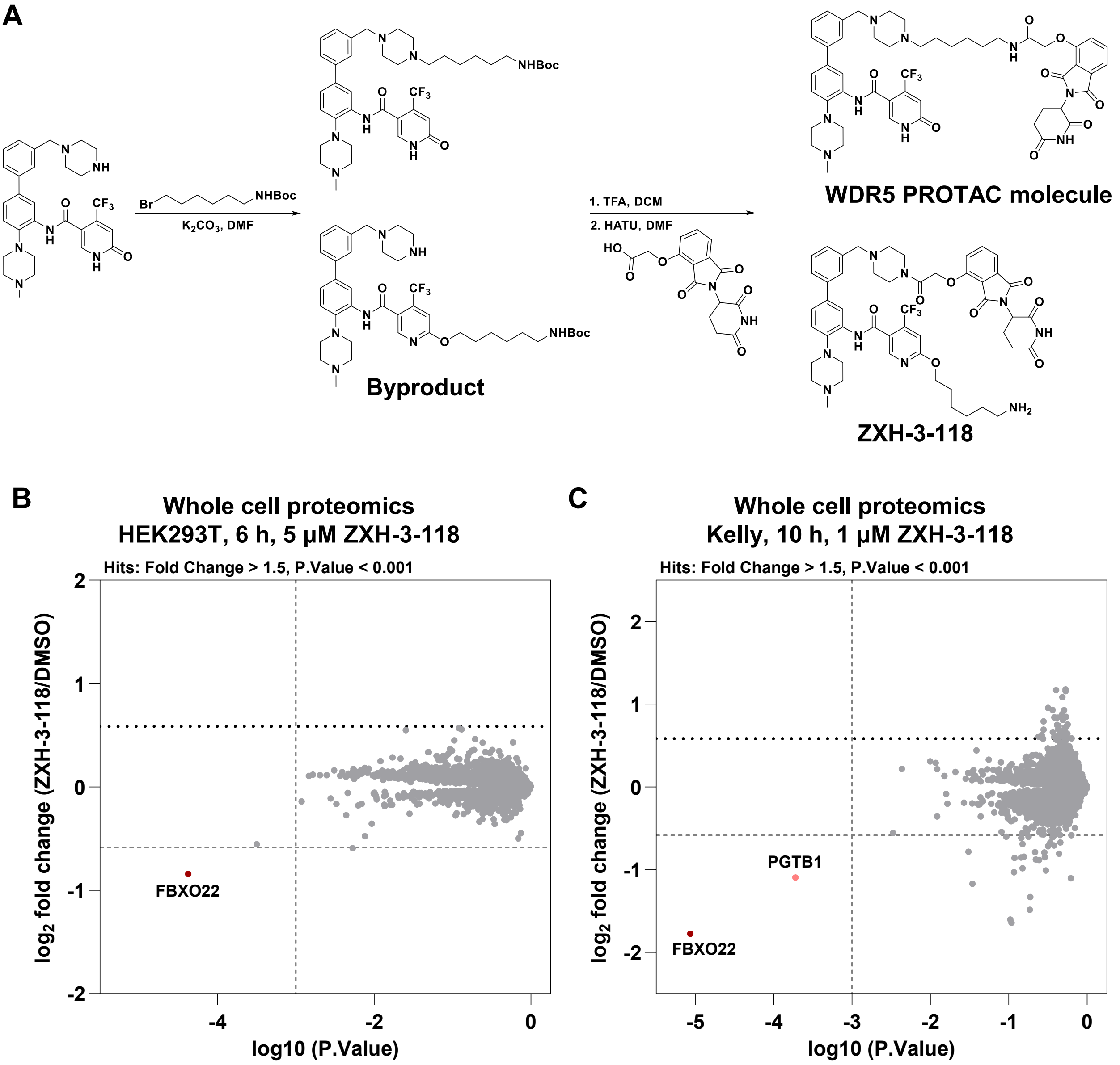


**Figure S1.** Chemo-proteomics reveal unexpected degradation of FBXO22 by a CRBN based degrader. A) Chemical synthesis scheme of ZXH-3-118. B) Quantitative proteome-wide mass spectrometry in HEK293T cells after 6 hours treatment with 5 µM ZXH-3-118. C) Quantitative proteome-wide mass spectrometry in Kelly cells after 10 hours treatment with 1 µM ZXH-3-118.

#### SI-2. ZXH-03-118 degrades FBXO22 in different cell lines

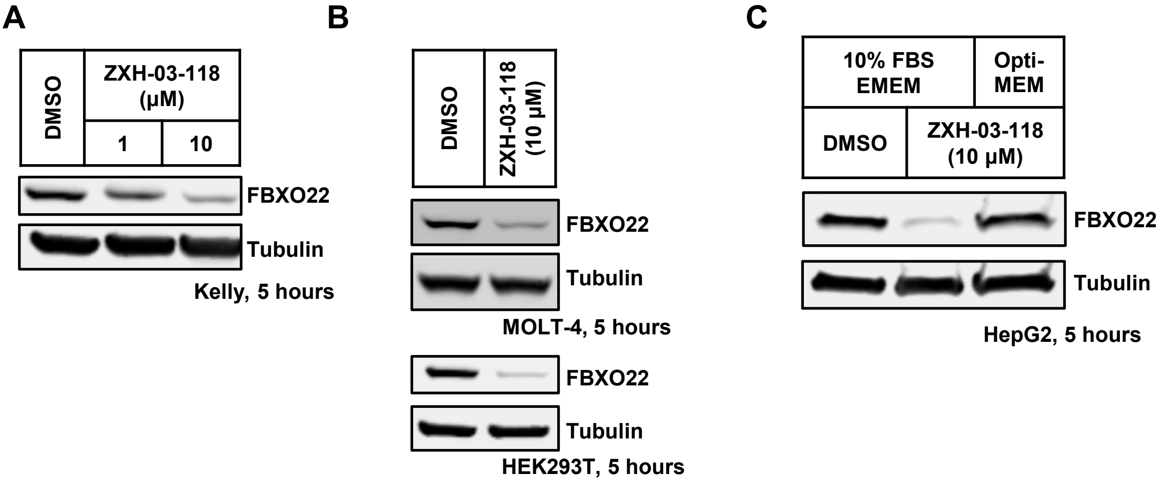


**Figure S2.** ZXH-03-118 degrades FBXO22 in different cell lines。 A) Western blots showing protein levels of FBXO22 in Kelly cells treated with indicated concentration of ZXH-03-118 for 5 hours. B) Western blots showing protein levels of FBXO22 in MOLT-4 or HEK293T cells treated with 10 μM ZXH-03-118 for 5 hours. C) Western blots showing protein levels of FBXO22 in HepG2 cells treated with 10 μM ZXH-03-118 in indicated media for 5 hours.

#### SI-3. Screening of CRBN based degrader compounds

**A**

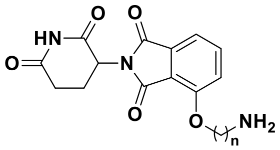


| n | $DC_{50}$ | $D_{Max}$ |
|---|---|---|
| 2,4,6,8 | > 10 uM | N.D. |

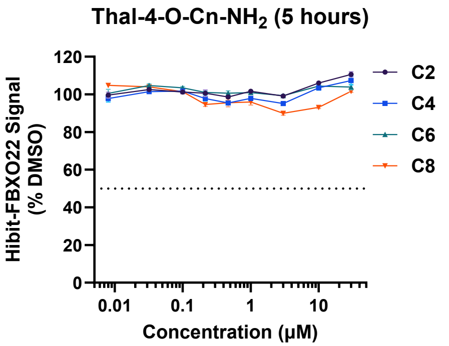


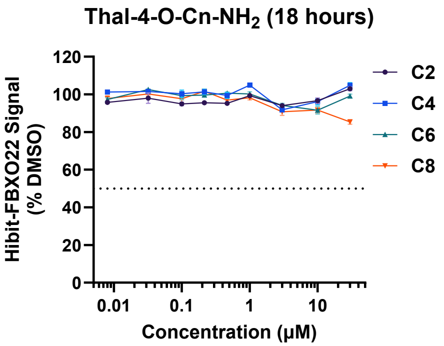


**B**

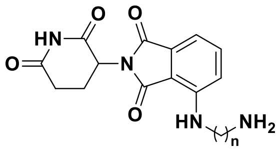


| n | $DC_{50}$ | $D_{Max}$ |
|---|---|---|
| 2/3/4/5/6/7 PEG2 | > 10 uM | N.D. |
| 8 | > 10 uM | 21% |

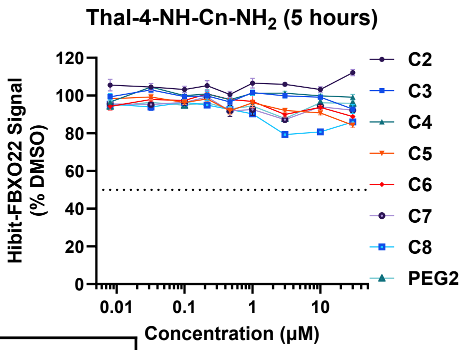


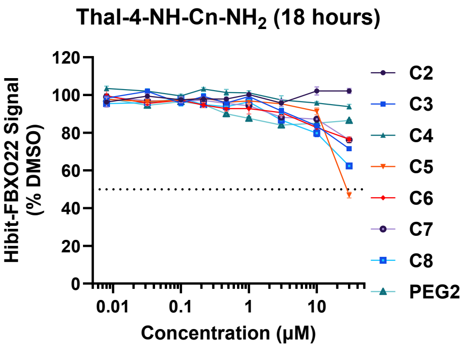


**C**

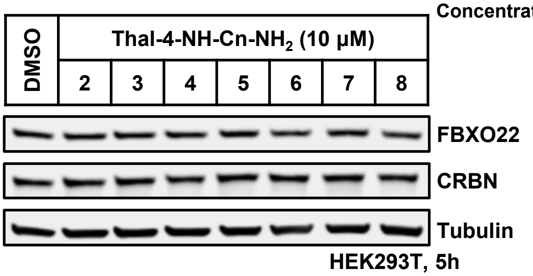

**D**

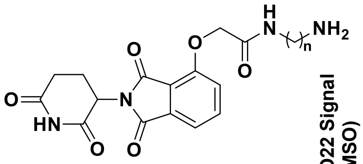


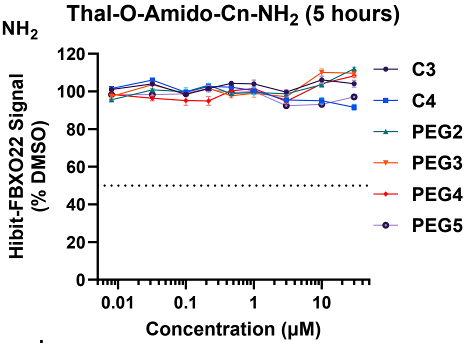


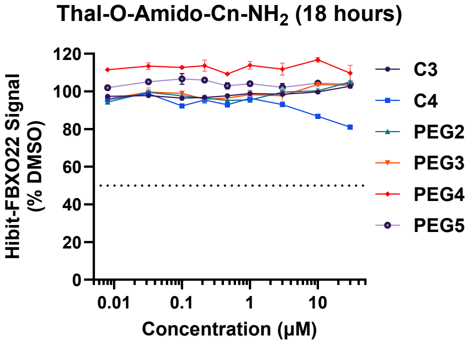


| | 5 h | | 18 h | |
|---|---|---|---|---|
| n | $DC_{50}$ | $D_{Max}$ | $DC_{50}$ | $D_{Max}$ |
| 3/5 | > 10 µM | N.D. | > 10 µM | N.D. |
| PEG2/3/4/5 | > 10 µM | N.D. | > 10 µM | N.D. |
| 4 | > 10 µM | N.D. | > 10 µM | 13% |
| 6 | > 10 µM | 42% | > 10 µM | 33% |
| 7 | > 10 µM | 32% | > 10 µM | 27% |
| 8 | 2.1 µM | 53% | > 10 µM | 37% |

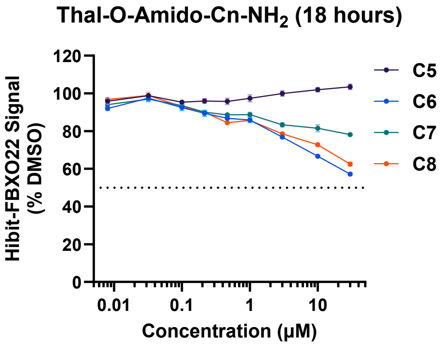


**Figure S3.** Screening of CRBN based degrader compounds. A) General chemical structure of Thalidomide-O-Cn-$NH_2$ and HiBiT-FBXO22 assay results for Jurkat cells treated with the CRBN ligand-based degraders for 5 or 18 hours. B) General chemical structure of Thalidomide-NH-Cn-$NH_2$ and HiBiT-FBXO22 assay results for Jurkat cells treated with the CRBN ligand-based degraders for 5 or 18 hours. C) Western blots showing protein levels of FBXO22 and CRBN in HEK293T cells treated with indicated compounds for 5 hours. D) General chemical structure of Thalidomide-Amide-Cn-$NH_2$ and HiBiT-FBXO22 assay results for Jurkat cells treated with the CRBN ligand-based degraders for 5 or 18 hours. Tables summarized the $DC_{50}$ and $D_{max}$ from Hibit-FBXO22 assays.

#### SI-4. Screening of VHL based degrader compounds

**A**

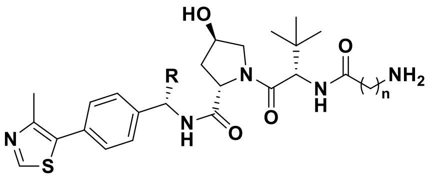


| | | 5 hours | | 18 hours | |
|---|---|---|---|---|---|
| **R** | **n** | **$DC_{50}$** | **$D_{Max}$** | **$DC_{50}$** | **$D_{Max}$** |
| H | 4/5 | > 10 μM | N.D. | > 10 μM | N.D. |
| H | 6 | 0.42 μM | 73% | 0.079 μM | 79% |
| **Me** | **6** | **0.29 μM** | **82%** | **0.055 μM** | **86%** |
| H | 7 | 1.7 μM | 55% | 0.34 μM | 66% |
| H | 10 | 0.29 μM | 71% | 0.71 μM | 87% |
| H | PEG2 | > 10 μM | 8% | > 10 μM | 35% |
| H | PEG3 | > 10 μM | 43% | 2.3 μM | 75% |
| H | PEG4 | > 10 μM | N.D. | > 10 μM | 8% |

**B**

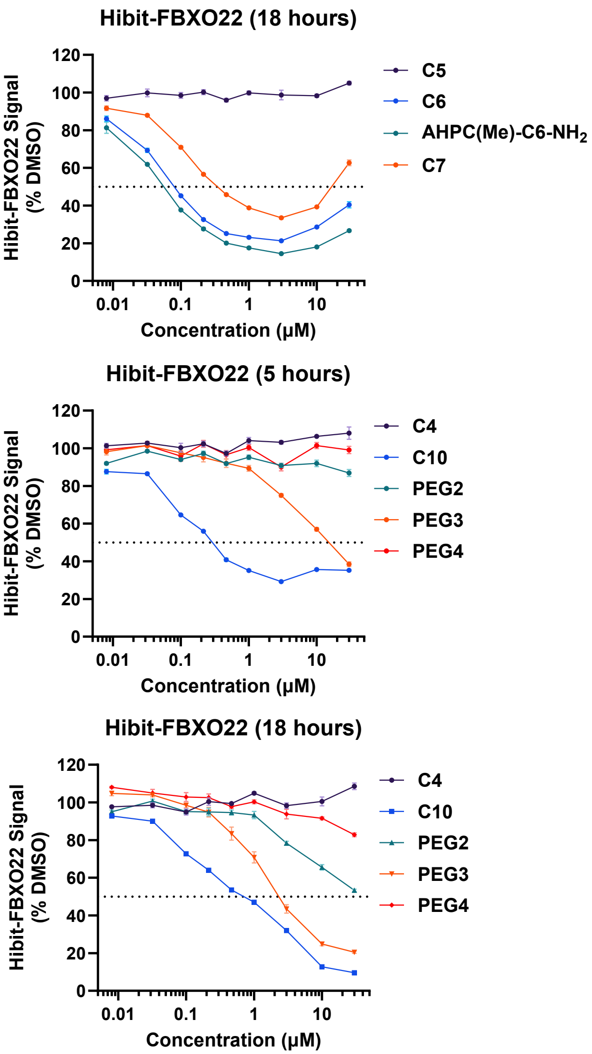


**Figure S4.** Screening of VHL based degrader compounds. (A) General chemical structure of AHPC(R)-Cn-$NH_2$. (B) HiBiT-FBXO22 assay results for Jurkat cells treated with the VHL ligand-based degraders for 5 or 18 hours. Tables summarized the $DC_{50}$ and $D_{max}$ from Hibit-FBXO22 assays.

#### SI-5. Screening of MDM2/IAP based degrader compounds

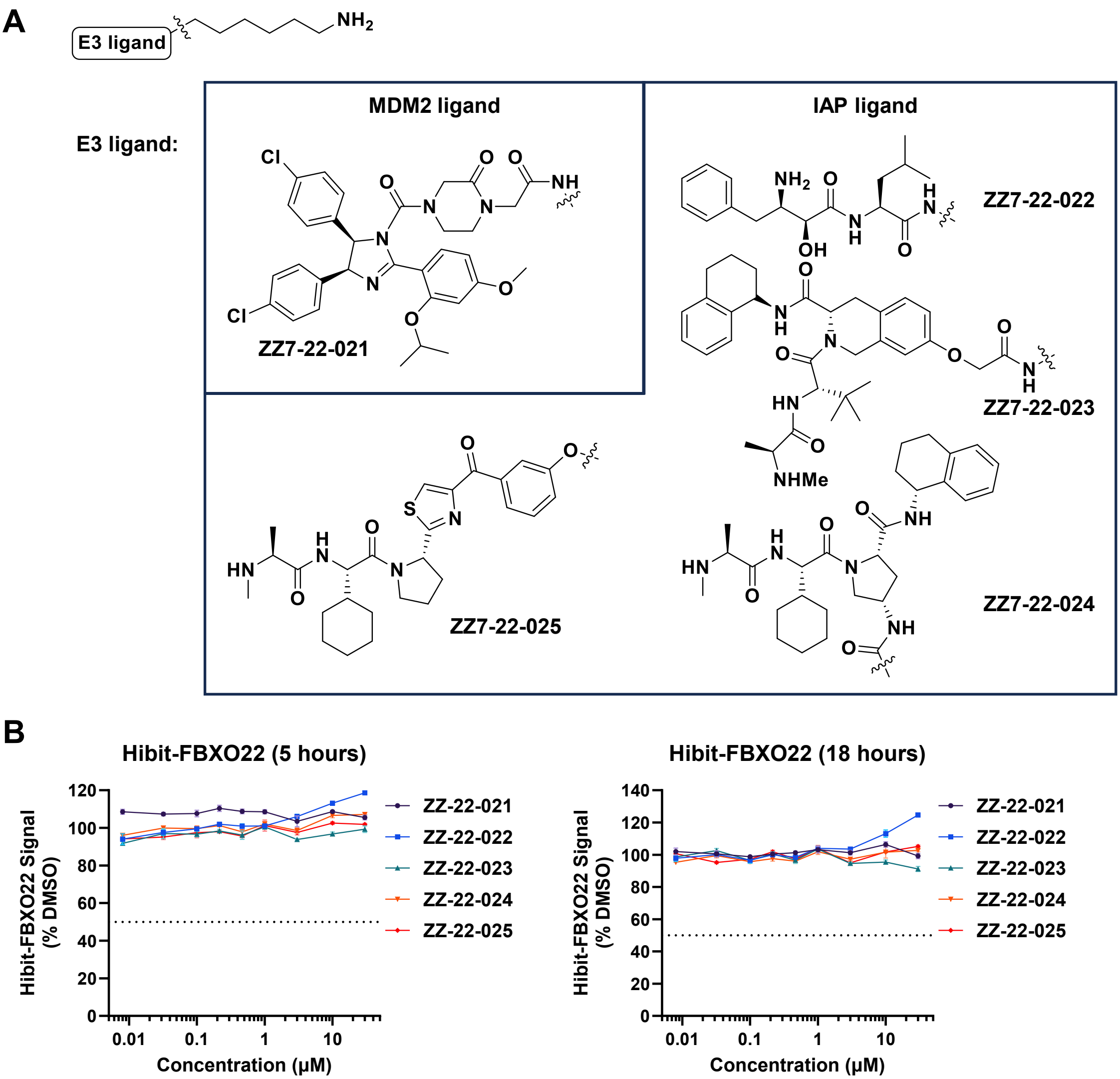


**Figure S5.** Screening of MDM2/IAP based degrader compounds. A) Chemical structure of primary amine tethered with MDM2 or IAP ligand. B) HiBiT-FBXO22 assay results for Jurkat cells treated with the MDM2 or IAP ligand-based degraders for 5 or 18 hours.

#### SI-6. Hibit assay to assess the roles of the UPS pathway and primary amine metabolic conversion

**A**

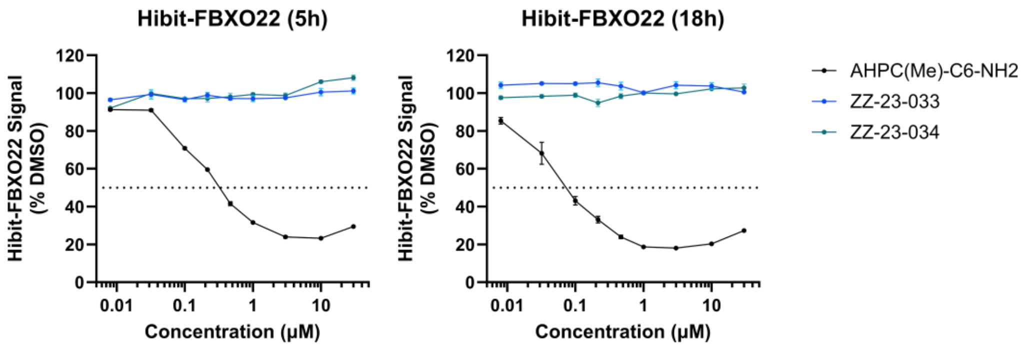


**B**

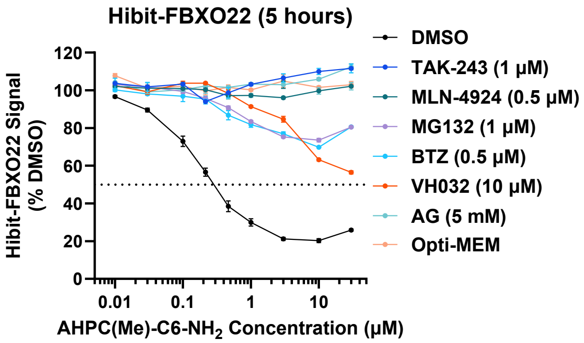


**Figure S6.** Hibit assay to assess the roles of the UPS pathway and primary amine metabolic conversion. A) HiBiT-FBXO22 assay results for Jurkat cells treated with the indicated compounds for 5 hours or 18 hours. B) HiBiT-FBXO22 assay results for Jurkat cells pre-treated with the indicated inhibitors for 1 hour, followed by treatment with 1 μM AHPC(Me)-C6-$NH_2$ for 5 hours. The “Opti-MEM” condition indicates treating the compound in Opti-MEM media.

#### SI-7. SAR on AHPC-Cn-CHO

**A**

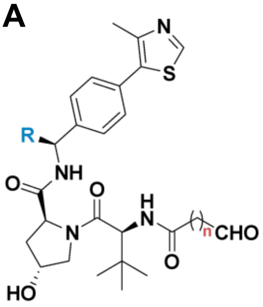


| | R Group | Linker (n) |
|---|---|---|
| ZZ7-24-008 | Me | 0 |
| TWQ-07-40 | Me | 4 |
| AHPC-CHO | H | 5 |
| TWQ-07-41 | Me | 6 |

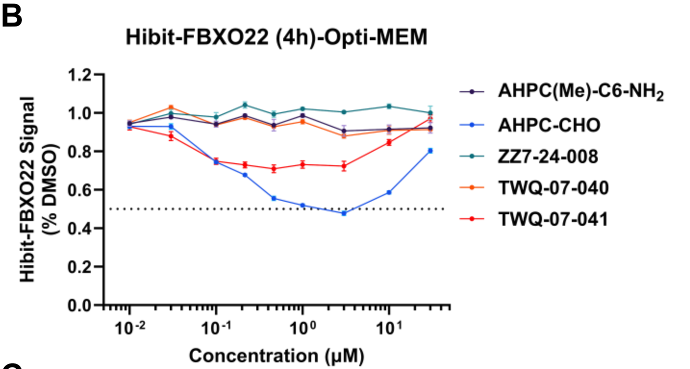


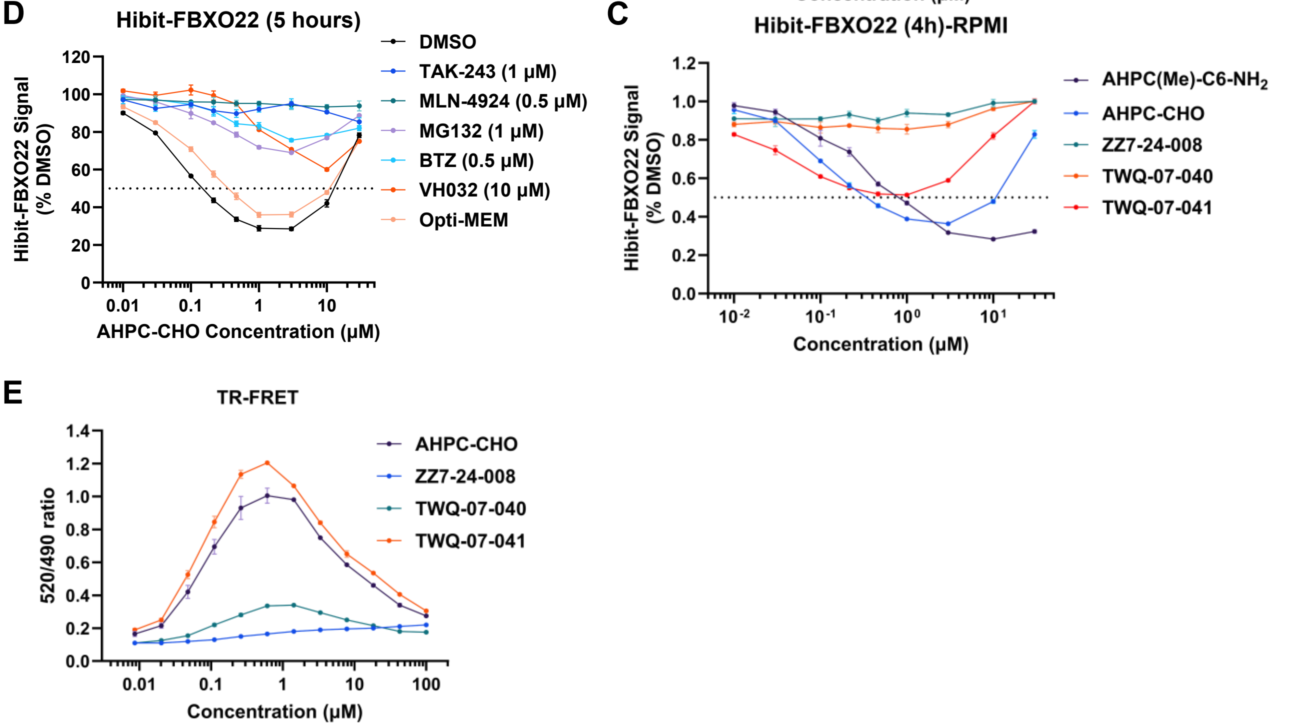


**Figure S7.** SAR on AHPC-Cn-CHO. A) General chemical structure of AHPC-Cn-CHO. B) HiBiT-FBXO22 assay results for Jurkat cells treated with the VHL ligand-based degraders in Opti-MEM media for 4 hours. C) HiBiT-FBXO22 assay results for Jurkat cells treated with the VHL ligand-based degraders in RPMI media for 4 hours. D) HiBiT-FBXO22 assay results for Jurkat cells pre-treated with the indicated inhibitors for 1 hour, followed by treatment with 1 μM AHPC-CHO for 5 hours. E) TR-FRET assay to measure molecule-dependent ternary complex formation between VCB complex and SKP1/FBXO22 with indicated compounds. Each point represents a duplicate replicates; the mean value line is drawn.

#### SI-8. Generation of FBXO22 KO Jurkat cell line

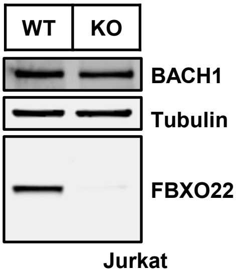


**Figure S8.** Western blots conform the knockout of FBXO22 in Jurkat cells.

### SI-9. Discovery of primary diamine compounds as FBXO22 self-degrader

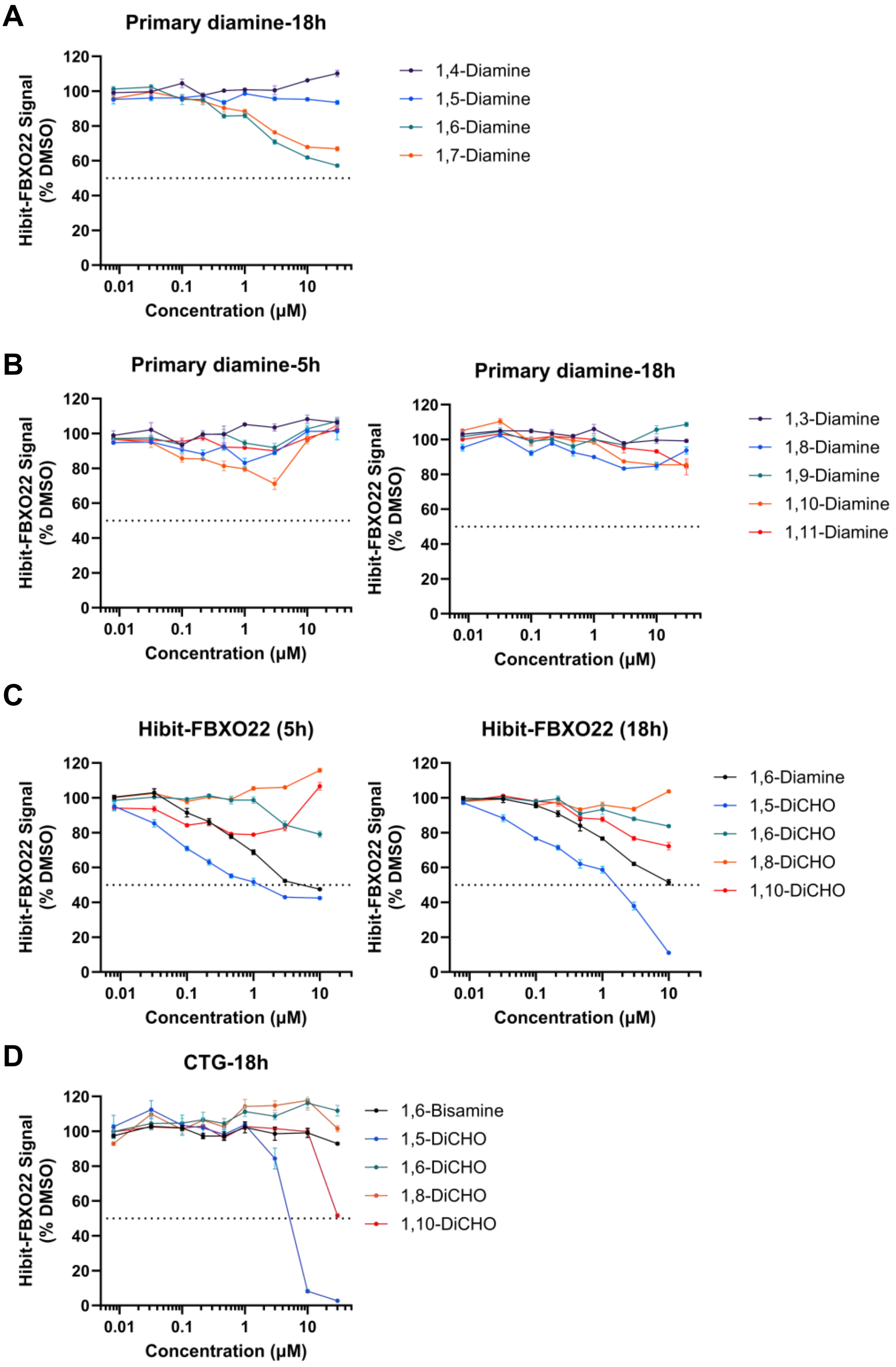


**Figure S9.** Discovery of primary diamine compounds as FBXO22 self-degrader. A) HiBiT-FBXO22 assay results for Jurkat cells treated with the primary diamines for 18 hours. B) HiBiT-FBXO22 assay results for Jurkat cells treated with the primary diamines for 5 or 18 hours. C) HiBiT-FBXO22 assay results for Jurkat cells treated with the dialdehydes for 5 or 18 hours. D) Cell viability assay results for Jurkat cells treated with indicated compounds for 18 hours.

#### SI-10. Alkyl amine or alkyl aldehyde is not a universal warhead for FBXO22 dependent protein degradation

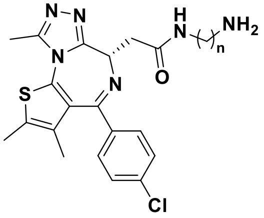


| | |
|---|---|
| n = 3 | JQ1-C3-$NH_2$ |
| n = 8 | JQ1-C8-$NH_2$ |

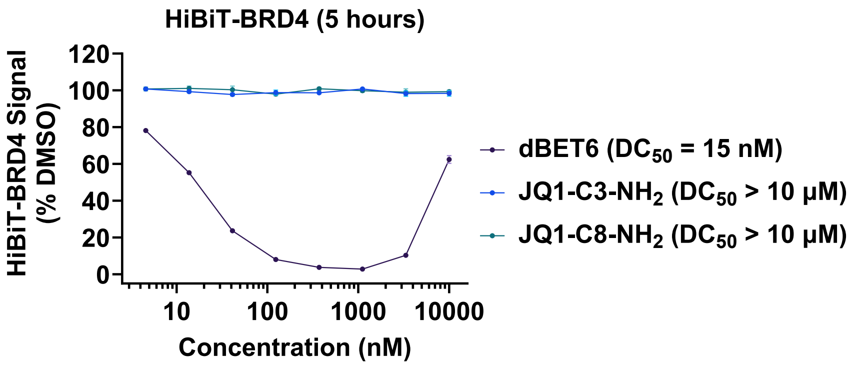


**Figure S10.** Alkyl amine or alkyl aldehyde is not a universal warhead for FBXO22 dependent protein degradation. General chemical structure of JQ1-Cn-$NH_2$ and HiBiT-BRD4 assay results for Jurkat cells treated with the indicated compounds for 5 hours. dBET6 is used as positive compound.

#### SI-11. Hibit and TR-FRET analysis of VHL ligand-Covalent warhead compounds

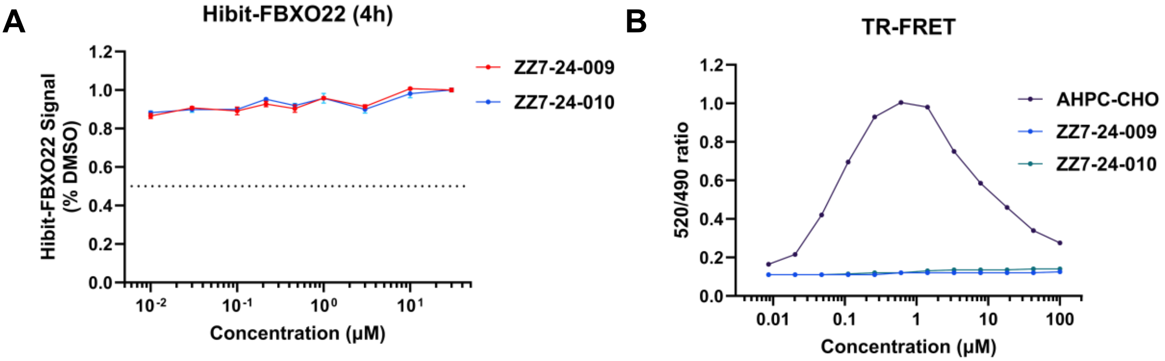


**Figure S11.** Hibit and TR-FRET analysis of VHL ligand-Covalent warhead compounds. A) HiBiT-FBXO22 assay results for Jurkat cells treated with the VHL ligand-based degraders for 4 hours. B) TR-FRET assay to measure molecule-dependent ternary complex formation between VCB complex and SKP1/FBXO22 with indicated compounds. Each point represents a duplicate replicates; the mean value line is drawn.

#### SI-12. Analysis of 2-PCA based FBXO22 degrader compounds

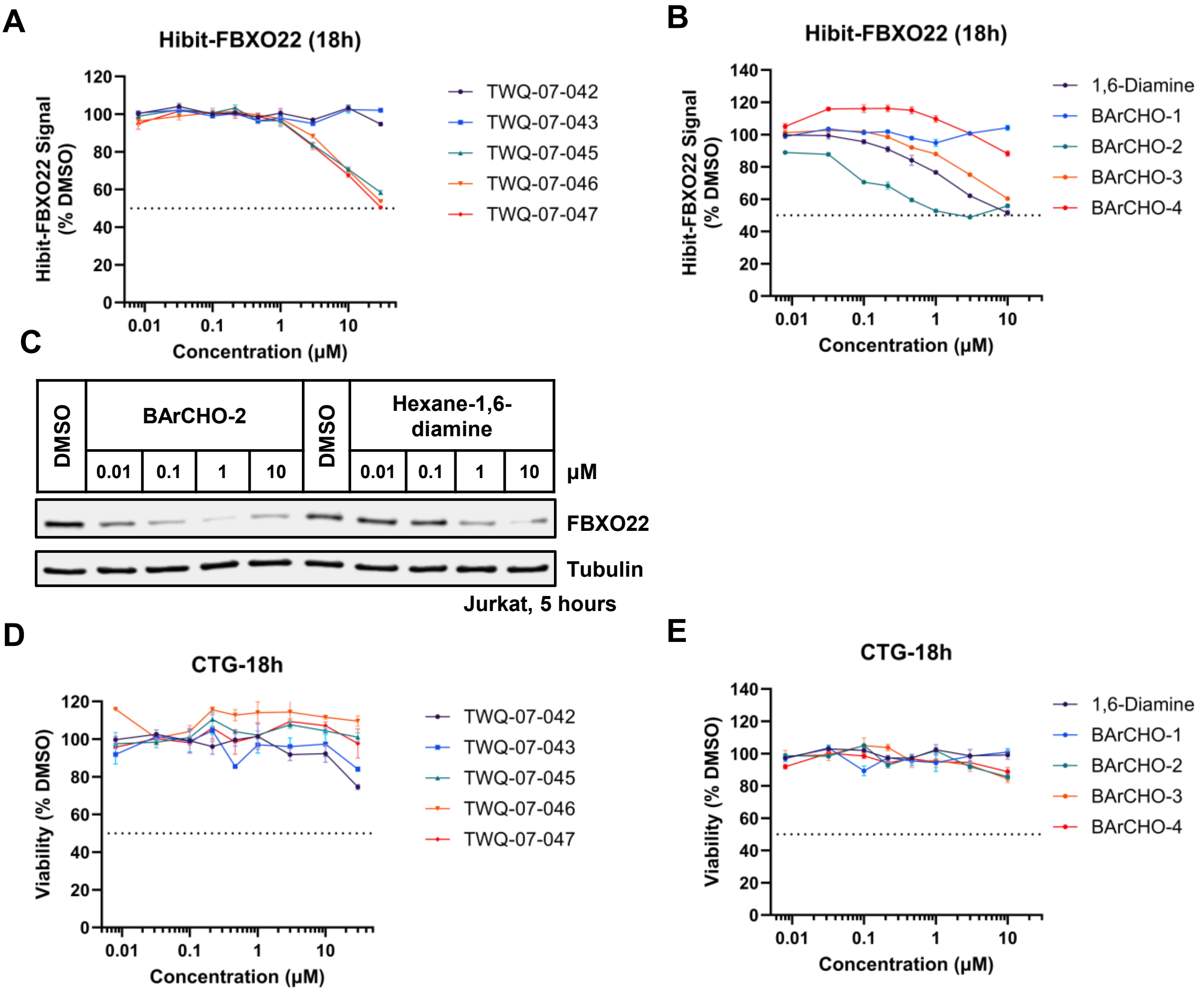


**Figure S12.** Analysis of 2-PCA based FBXO22 degrader compounds. A) HiBiT-FBXO22 assay results for Jurkat cells treated with the indicated compounds for 5 hours. B) HiBiT-FBXO22 assay results for Jurkat cells treated with the indicated compounds for 18 hours. C) Western blots showing protein levels of FBXO22 in Jurkat cells treated with indicated concentration of BArCHO-2 and Hexane-1,6-diamine for 5 hours. D) Cell viability assay results for Jurkat cells treated with indicated compounds for 18 hours. E) Cell viability assay results for Jurkat cells treated with indicated BArCHOs for 18 hours.

#### SI-13. Linker SAR on JQ1-PCA

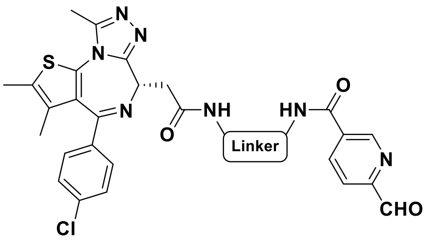


| Linker | Compound ID |
|---|---|
| C4 | ZZ7-18-064 |
| C6 | ZZ7-18-065 |
| C8 | ZZ7-18-066 |
| PEG2 | ZZ7-18-068 |
| PEG4 | ZZ7-18-069 |

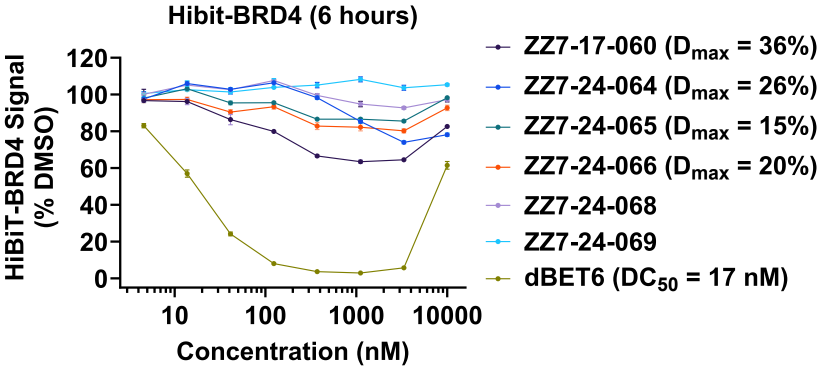


**Figure S13.** Linker SAR on JQ1-PCA. General chemical structure of JQ1-PCA and HiBiT-BRD4 assay results for Jurkat cells treated with the indicated compounds for 6 hours. dBET6 is used as positive compound.

#### SI-14. SAR on JQ1-2PCA

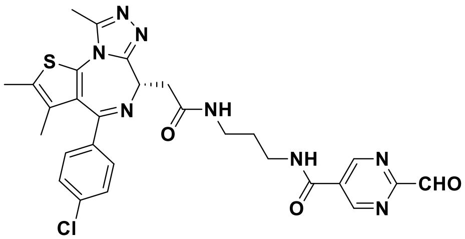

ZZ7-24-030

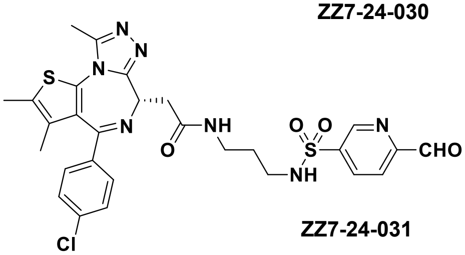

ZZ7-24-031

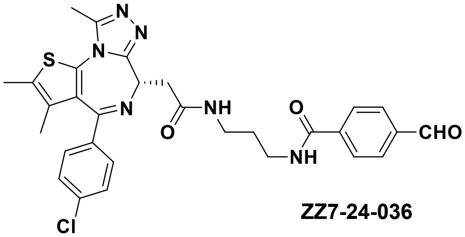

ZZ7-24-036

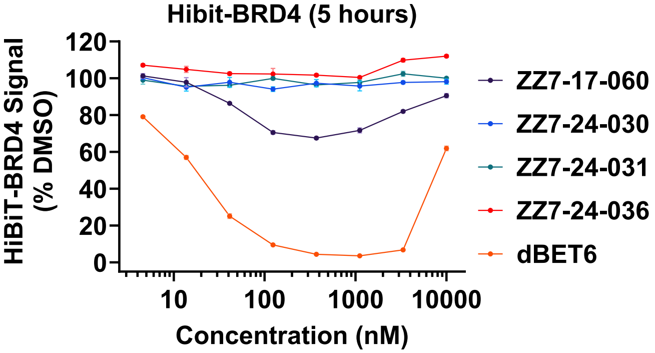


**Figure S14.** Warhead SAR on JQ1-2PCA. Chemical structure of aromatic aldehyde tethered JQ1 and HiBiT-BRD4 assay results for Jurkat cells treated with the indicated compounds for 5 hours. dBET6 is used as positive compound.

**SI-15. Ubiquitin-proteasome system (UPS)-focused CRISPR screen for $BRD4_{BD1}$-eGFP stability in K562-Cas9 cells treated with 10 µM ZZ7-17-060 for 16 hours.**

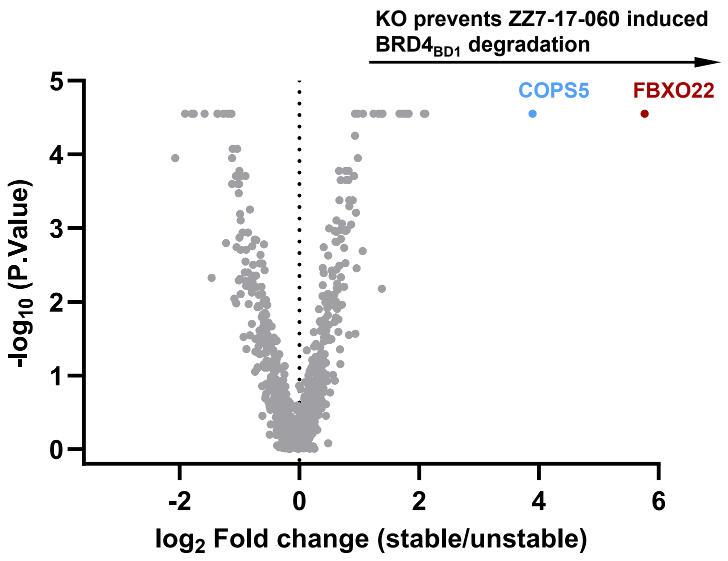


**Figure S15.** Ubiquitin-proteasome system (UPS)-focused CRISPR screen for $BRD4_{BD1}$-eGFP stability in K562-Cas9 cells treated with 10 µM ZZ7-17-060 for 16 hours. Note the adaptor protein SKP1 is not in the BISON library.

#### SI-16. FBXO22 KO completely rescue ZZ7-17-060 induced BRD4 degradation

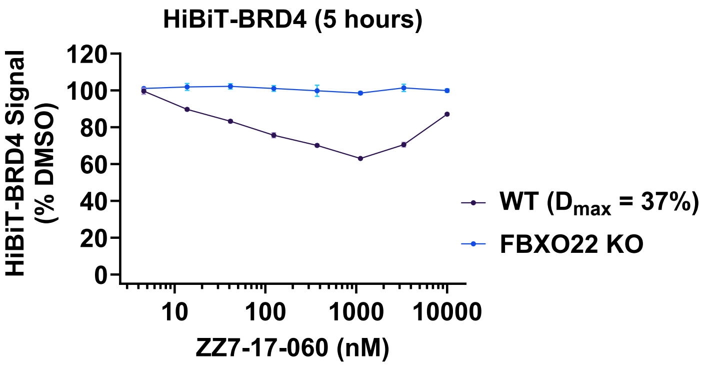


**Figure S16.** FBXO22 KO completely rescue ZZ7-17-060 induced BRD4 degradation. HiBiT-BRD4 assay results in WT Jurkat cells or FBXO22 KO Jurkat cells treated with the indicated compounds for 5 hours.

#### SI-17. Reintroducing WT FBXO22 but not C326A recapitulate ZZ7-17-060 mediated BRD4 degradation

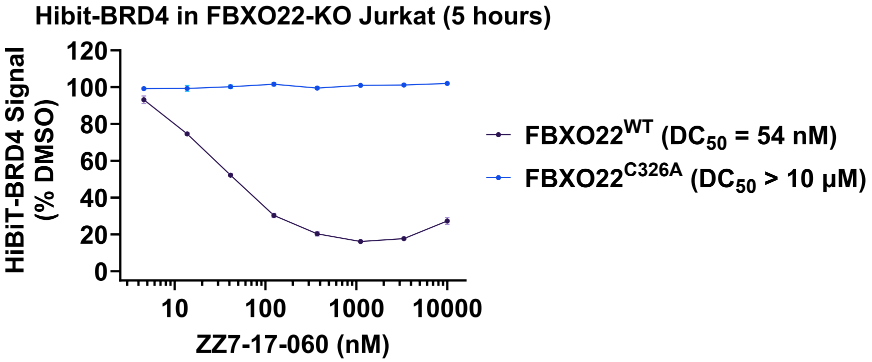


**Figure S17.** Reintroducing WT FBXO22 but not C326A recapitulate ZZ7-17-060 mediated BRD4 degradation. HiBiT-BRD4 assay results in re-overexpression WT FBXO22 or C326A FBXO22 in FBXO22 KO Jurkat cells treated with the indicated compounds for 5 hours.
